# Heterogeneous plasticity rules generate diverse structures in recurrent networks

**DOI:** 10.64898/2026.10.26.760407

**Authors:** Claudia Cusseddu, Dylan Festa, Christoph Miehl, Julijana Gjorgjieva

## Abstract

How do local changes at individual synapses give rise to structured recurrent connectivity? Experiments have revealed diverse forms of spike-timing-dependent plasticity (STDP) across the brain, making it difficult to predict which circuit structures arise from different rules. We developed an analytical framework mapping excitatory Hebbian, anti-Hebbian, and symmetric STDP rules to directed three-neuron motifs. Individual homogeneous rules generate restricted subsets of motifs, and no single rule produces all motifs. Combining different rules generates motifs inaccessible to individual rules and yields the full motif repertoire. These predictions generalize to larger recurrent networks. The mapping is nevertheless highly non-unique, with distinct rules and parameters producing similar motif distributions, including distributions compatible with experimental connectivity. Thus, circuit structure constrains candidate plasticity rules but does not uniquely identify them. Finally, networks with similar motif distributions can nonetheless differ in their firing rates and pairwise correlations, suggesting that activity may provide an additional constraint on candidate plasticity rules.

## Introduction

During development and learning, neural circuits acquire and refine structured patterns of connectivity that support diverse types of computation^1–4^. Circuit organization reflects both molecular mechanisms that establish cellular and synaptic specificity and activity-dependent mechanisms that modify connectivity through experience^5–7^. Such activity-dependent changes allow neural circuits to adapt during development, learning, and memory and can involve both changes in synaptic strength and structural remodeling of synaptic connections^2,8–11^. We therefore ask how much of the specialized connectivity observed in recurrent circuits can arise from local activity-dependent mechanisms, particularly when structure is not already imposed by the initial connectivity or external input.

A prominent activity-dependent mechanism for generating structured connectivity is spike-timing-dependent plasticity (STDP), by which synapses change according to the relative timing of pre- and postsynaptic spikes. Early experimental characterizations of STDP in cortical circuits^12–14^ motivated the classical asymmetric Hebbian description, consistent with Hebb’s postulate that synapses are strengthened when presynaptic activity contributes to postsynaptic firing^15^. In its canonical form, presynaptic spikes preceding postsynaptic spikes induce long-term potentiation (LTP), whereas the reverse order induces long-term depression (LTD)^16,17^. This form of STDP has been studied extensively in theoretical models of recurrent connectivity^18–20^. Much of this work considered prespecified plasticity rules and asked which structures they generate. Yet experiments have revealed a much broader diversity of STDP rules that varies across cell types, brain regions, neuromodulatory states, and synapse classes^21–26^. In addition to asymmetric Hebbian rules, temporally asymmetric anti-Hebbian and temporally symmetric rules have been observed. Recent theoretical work has shown that changing the temporal causality of STDP can qualitatively alter recurrent structure^27^.

This diversity raises a central question: which aspects of recurrent circuit structure are constrained by a given plasticity rule, and how do these constraints change across different forms and parameters of STDP? The inverse question is equally important. If distinct plasticity rules can generate similar patterns of connectivity, to what extent can an observed circuit structure constrain the rules that produced it? Addressing these questions requires comparing different forms of STDP directly rather than studying individual rules in isolation.

These comparisons require a representation of circuit structure that can be used across different plasticity rules. Directly comparing full recurrent connectivity matrices is difficult because the number of possible wiring patterns grows combinatorially with network size. Local connectivity motifs provide a tractable alternative by describing small subnetworks that capture recurring patterns of circuit architecture^28–30^. Triplet motifs are particularly useful because they extend beyond pairwise connectivity while remaining analytically and experimentally accessible. Across species and brain regions, experiments have found distributions of local motifs that deviate from random expectations^31–36^. Different brain areas also exhibit distinct patterns of motif over- and under-representation^37^, suggesting that motif statistics capture meaningful differences in recurrent organization. The arrangement of motifs can also influence network dynamics, including synchronous covariability, dimensionality of activity, and the propagation of spiking patterns^38–43^. Related theoretical work has shown how the structure of local recurrent interactions can shape the network-wide effects of local perturbations ^44^. Triplet motifs therefore provide an intermediate scale for relating local plasticity rules to recurrent structure and for asking how much additional information is provided by network activity.

To establish this link, we develop an analytical framework that maps a broad range of excitatory STDP rules to directed triplet motifs. We parameterize Hebbian, anti-Hebbian, and symmetric rules through the balance between LTD and LTP and their relative time constants, and derive interaction coefficients that determine the resulting synaptic dynamics^45^. We consider networks receiving unstructured external input so that connectivity structure is not prescribed by patterned activity. We find that individual homogeneous rules realize only part of the possible motif space. Combining different rules allows a broader range of triplet structures to emerge. At the same time, the mapping from plasticity to structure is highly degenerate, so experimentally observed motif distributions narrow the range of candidate rules without specifying a unique mechanism. Finally, network activity can provide additional constraints when structurally similar networks arise from different plasticity mechanisms.

## Results

### Establishing a mapping between STDP rules and triplet motifs

We first asked how the form of an STDP rule constrains the local connectivity patterns that emerge in recurrent networks. Because the number of possible connectivity patterns grows rapidly with network size, we considered triplet connectivity motifs. These three-neuron circuits are the smallest subnetworks that go beyond pairwise interactions while remaining analytically tractable, making them a natural “building block” for linking synaptic plasticity rules to circuit architecture (Fig. 1A, B).

**Figure 1:**
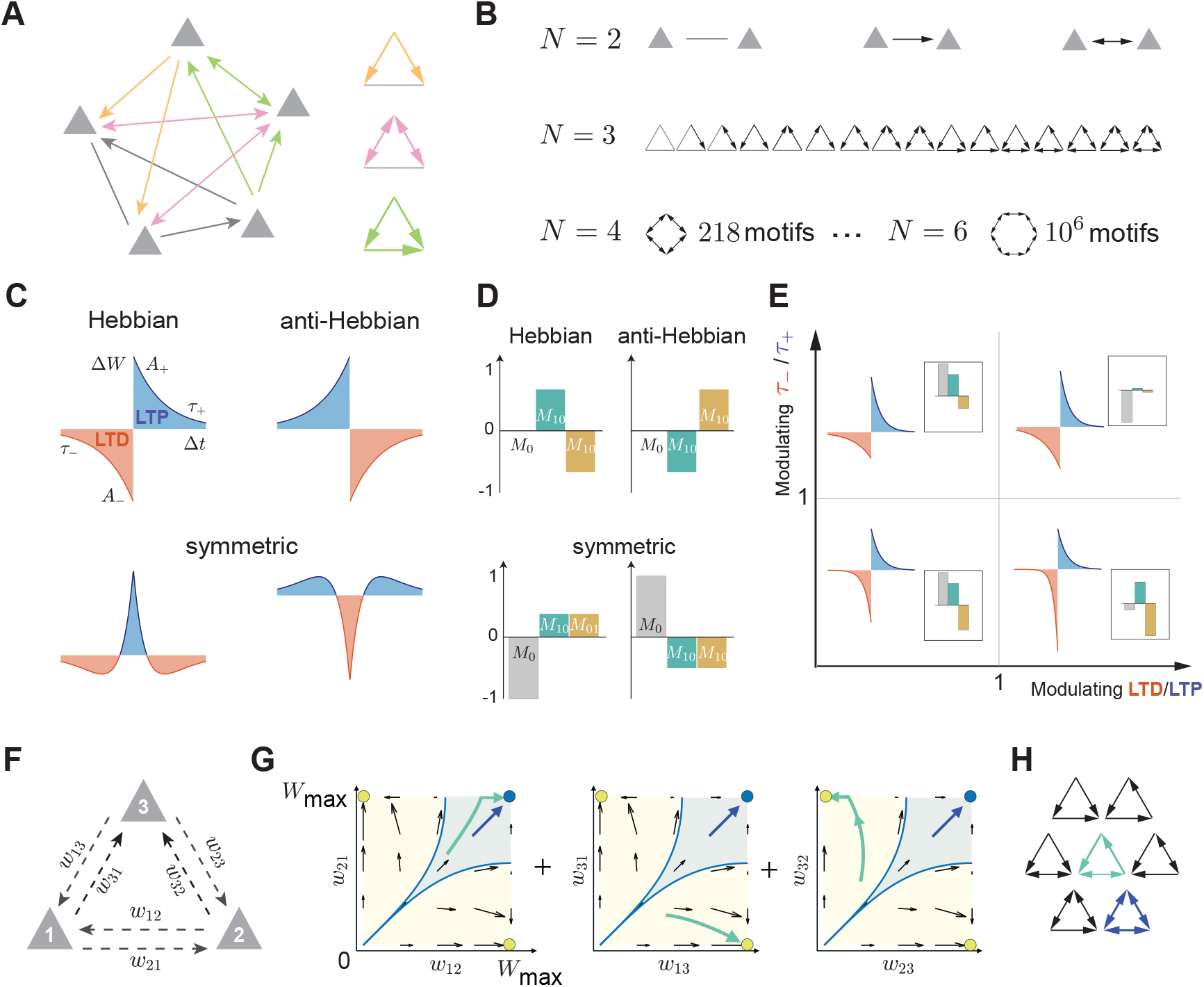
Parametrization of spike-timing-dependent plasticity (STDP) rules and emergence of triplet motifs. **A.** *Left*: Schematic of a recurrent circuit. *Right*: Examples of triplet motifs embedded in the network. **B**. Number of possible motifs as the number of neurons (*N*) increases from *N* = 2 to *N* = 6. **C**. Parametrized STDP rules grouped into Hebbian, anti-Hebbian, and symmetric families. Blue and red areas indicate total long-term potentiation (LTP) and long-term depression (LTD), respectively. Synaptic change Δ*W* is shown as a function of the spike time difference Δ*t* = *t*_post_ − *t*_pre_. *A*_+_ and *A*_−_ denote the maximum amplitudes of LTP and LTD, and *τ*_+_ and *τ*_−_ their respective decay time constants. **D**. Motif coefficients for example STDP rules (cf. Eq. 1). *Top*: Hebbian and anti-Hebbian rules, both with LTD = LTP, *τ*_−_ = *τ*_+_. *Bottom*: Symmetric rules with LTD = 1.05× LTP and *τ*_−_ = 3 *τ*_+_ and their inverse values, respectively. **E**. Examples of how the STDP rule and motif coefficients (insets) change with the ratio of time constants, *τ*_−_/*τ*_+_, and the ratio of total LTD and LTP areas. **F**. Three-neuron circuit with reciprocal connections *w*_*ij*_ and *w*_*ji*_, where *i, j* ∈ {1, 2, 3}, before plasticity. **G**. Phase planes for the three pairs of synaptic weights under a Hebbian STDP rule with LTD/LTP = 1 and *τ*_−_/*τ*_+_ = 2. Weights are bounded within [0, *w*_max_]. Green and blue lines show example trajectories, shaded regions indicate their basins of attraction, and colored circles mark the corresponding attractors. **H**. All possible steady-state triplet motifs arising from the dynamics shown in G. Colors correspond to the two example trajectories in **G**.

To relate spiking activity to synaptic change, we used a framework based on multivariate mutually excitatory Hawkes processes^46^ (Methods). Within this framework, synaptic change can be expressed analytically in terms of firing rates, spike train correlations, and network connectivity. Assuming a separation of timescales between fast spiking and slow plasticity, the expected change of a synapse *w*_*ij*_ from neuron *i* to neuron *i* can be approximated by^45,47–50^:

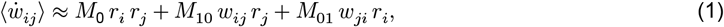

with hard bounds 0 ≤ *w*_*ij*_ ≤ *w*_max_, where *r*_*j*_ and *r*_*i*_ are the pre- and postsynaptic firing rates, respectively.

Eq. (1) separates synaptic change into three contributions. The coefficient *M*_0_ describes the contribution from independent spiking, *M*_10_ multiplies the direct feedforward connection for the neuron pair, *w*_*ij*_, and *M*_01_ multiplies the reciprocal connection *w*_*ji*_. Their signs and relative magnitudes determine whether a pair of synapses converges to absent, unidirectional, or bidirectional connectivity. These coefficients are determined by the STDP rule and the excitatory postsynaptic potential (EPSP) function rather than chosen independently (Methods; Fig. S1). When firing rates are kept approximately constant over the plasticity timescale, the STDP rule therefore determines the synaptic weight dynamics and the resulting motif.

Because experimentally measured STDP profiles vary widely^21^, their corresponding motif coefficients should also vary across rules. We therefore asked which triplet motifs can emerge at steady state for a given STDP rule and which cannot emerge under that rule. To address this question, we established a rule-to-motif mapping from a broad class of STDP rules to their resulting triplet motifs.

We grouped experimentally measured STDP rules into three families: Hebbian, anti-Hebbian, and symmetric rules (Fig. 1C). We parameterized each family by the relative strength of LTD and LTP and the ratio of their characteristic time constants, so that each coefficient in Eq. 1 could then be expressed as a function of the rule shape, *M*_•_ = *f* (LTD/LTP, *τ*_−_/*τ*_+_) (Fig. 1D, E). This allowed us to determine how changes in the STDP parameters alter the phase-plane dynamics of each synapse pair and, consequently, the resulting triplet motifs (Fig. 1F-H, Fig. S2). Synaptic weights were bounded between zero and *w*_max_, such that attractors at the lower and upper bounds correspond to absent and strengthened connections, respectively. A fixed rule can support several attractors with distinct basins of attraction for a connection pair, while varying the rule parameters shifts the system between regimes that support different motif classes.

### Hebbian rules produce a subset of triplet motifs

We analyzed the steady-state triplet motifs generated by Hebbian rules, first measured experimentally^12^, as a function of LTD/LTP and *τ*_−_/*τ*_+_ (Fig. 2). When the firing rates of the three neurons were kept constant, the two ratios alone determined the plasticity of the connections. We identified five qualitatively distinct motif classes, characterized by the loss of connections, the formation of unidirectional or bidirectional connections, or the coexistence of several outcomes depending on the initial conditions (Fig. 2A). When LTD dominated, all connections were eliminated and the three-neuron circuit converged to a disconnected state. In contrast, sufficiently strong LTP produced a fully connected state in which both weights of a bidirectional connection saturated at the upper bound, *w*_max_. Between these regimes, the outcome depended on the initial conditions, yielding motifs that combined absent and unidirectional connections, or unidirectional and bidirectional connections. Thus, Hebbian rules generated several motif classes but not the complete repertoire. In particular, no parameter regime produced motifs containing both bidirectional and absent connections.

**Figure 2:**
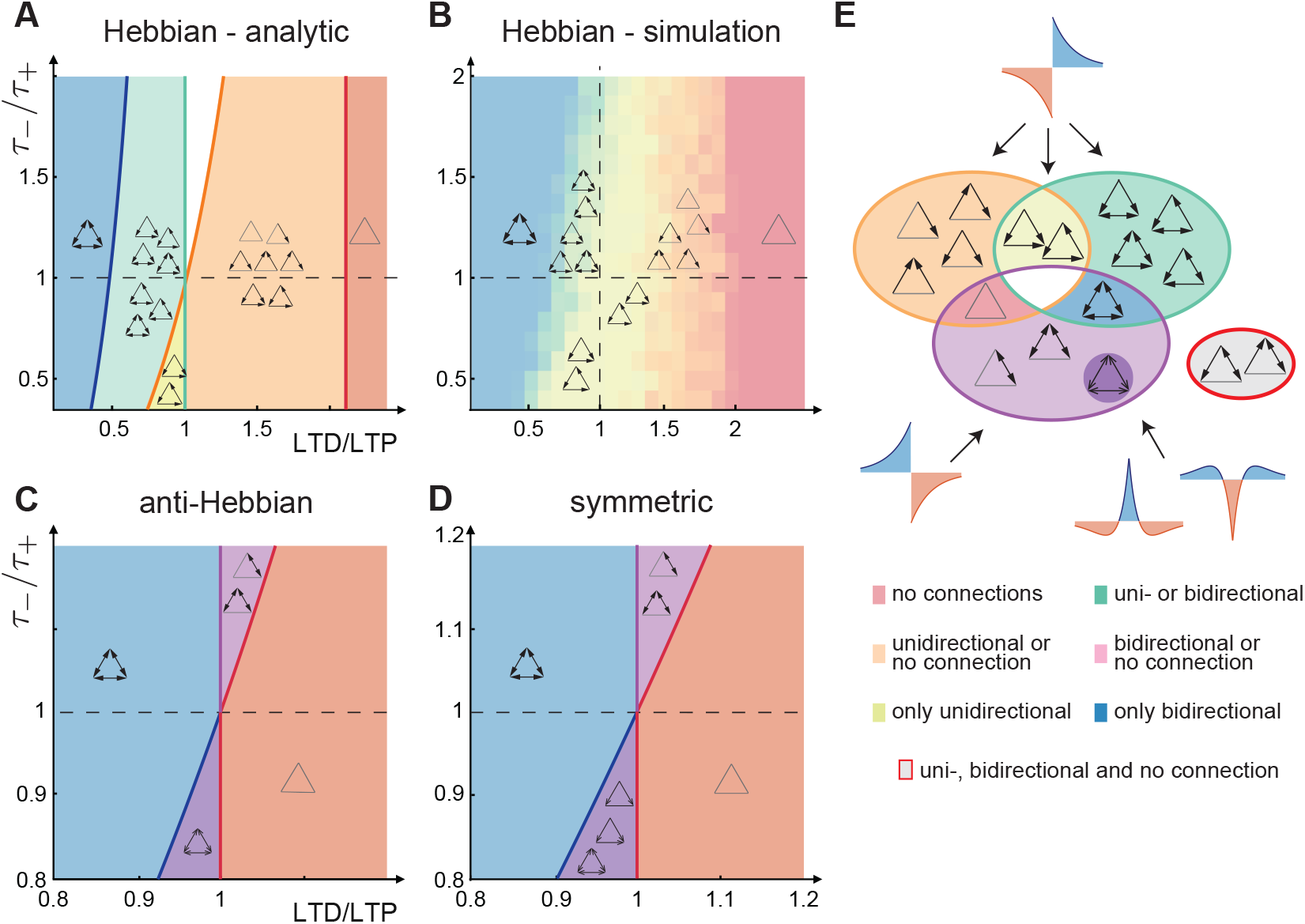
Emergence of triplet motifs under different STDP rules. **A.** Analytical prediction of triplet motifs generated by Hebbian rules as a function of LTD/LTP and *τ*_−_/*τ*_+_ (Supplementary Information S1). Colors indicate motif classes: no connections (red), absent or unidirectional connections (orange), only unidirectional connections (yellow), unidirectional or bidirectional connections (green), and only bidirectional connections (blue). In the analytically undetermined white region, all motifs are possible in principle. **B**. Motif classes obtained from simulations of three-neuron circuits with Hawkes dynamics under Hebbian rules, averaged over 10 trials for each parameter combination. The simulations support only unidirectional connections (yellow configuration) in the white region of A. **C**. Analytical prediction for anti-Hebbian rules as a function of LTD/LTP and *τ*_−_/*τ*_+_. Colors indicate no connections (red), only bidirectional connections (blue), absent or bidirectional connections (light purple), and bidirectional connections with weights spanning a continuous range (dark purple). **D**. Analytical prediction for symmetric rules as a function of LTD/LTP and *τ*_−_/*τ*_+_. Colors as in C, except for the dark purple region, where synaptic weights converge to a line attractor spanning a continuous range of unidirectional and bidirectional connections, which allows additional motifs (see Fig. S3). For panels A-D, the precise boundaries depend on the model parameters (Supplementary Information S1). Here, *w*_max_ = 0.5, *r* = 15 Hz, *τ* = 10 ms, *τ*_+_ = 20 ms, and *C* = 1. **E**. Summary of the motifs generated by the three rule families. Colors correspond to the classes in A-D. The gray group outlined in red contains motifs that cannot be generated by any of these individual rules.

Although the balance between LTD and LTP was the main determinant of motif class, their relative time constants could also change the outcome. In the yellow region of Fig. 2A, LTP slightly dominated over LTD in total area, yet only unidirectional connections emerged. Here, *τ*_+_ > *τ*_−_, such that LTD acted over a shorter timescale and counteracted potentiation for closely spaced post-pre spike pairs, preventing the formation of bidirectional connections. Conversely, when the total LTD and LTP areas were balanced, bidirectional connections emerged if *τ*_−_ was sufficiently larger than *τ*_+_. In this regime, LTP was concentrated within a shorter time window and could overcome the more broadly distributed LTD.

We validated our analytical predictions using simulations with three spiking neurons modeled as Hawkes processes (Fig. 2B), each paired with an inhibitory neuron that kept excitatory firing activity near a homeostatic target rate, through inhibitory plasticity^51^. All neurons received external, unstructured Poisson input spike trains at a fixed rate, so that connectivity structure was not imposed by the external drive but emerged from the interaction between the plasticity rule and the recurrent dynamics. The undetermined (white) region in the analytical prediction was particularly sensitive to the neuronal dynamics; while all motifs were possible in principle, the spiking dynamics favored configurations in which every neuron pair remained connected, either unidirectionally or bidirectionally. As a result, the simulations predominantly fell in the yellow and green regimes (Fig. 2B, yel-low and green). The simulations therefore confirmed how motif classes depend on both the LTD/LTP balance and their relative time constants. The rule-to-motif mapping between STDP rules and triplet motifs reveals how continuous variation in STDP parameters translates into discrete classes of circuit architecture.

### Anti-Hebbian and symmetric rules produce restricted but similar motif distributions

We next turned to anti-Hebbian and symmetric STDP rules to ask whether these rule families could produce the motifs that remained inaccessible under Hebbian STDP. Unlike Hebbian rules, anti-Hebbian and symmetric rules produced a more restricted set of triplet motifs (Fig. 2C,D). The two rule families produced largely overlapping motif classes, despite differences in their synaptic dynamics (Fig. S3). This overlap provides an example of the broad degeneracy in the rule-to-motif mapping as discussed below. The motifs produced by these two rule families were nonetheless complementary to those produced by Hebbian rules: anti-Hebbian and symmetric rules generated motifs that combined bidirectional coupling between one neuron pair with an absence of connections between the remaining pairs, including motifs that were inaccessible under Hebbian plasticity.

Furthermore, while Hebbian rules typically drove synapses toward the lower or upper bounds, 0 or *w*_max_ respectively, anti-Hebbian and symmetric rules could stabilize synaptic strengths across a continuous range, 0 ≤ *w*_*ij*_ ≤ *w*_max_, in specific parameter regimes. Overall, both families mainly led either to the loss of connections or to the formation of bidirectional connections. Under anti-Hebbian rules, both weights of a bidirectional connection could take values continuously within the bounds, and unidirectional connections never emerged. Symmetric rules additionally supported a region in which the synaptic weights converged to a line attractor, allowing unidirectional connections together with bidirectional ones across a continuous range (dark purple region in Fig. 2D; Fig. S3).

We confirmed these analytical predictions in simulations of three spiking neurons using the same Hawkes dynamics and homeostatic inhibition as for Hebbian rules (Fig. S3). Different plasticity families therefore impose distinct constraints on connectivity, and no single family produces the full spectrum of triplet motifs (Fig. 2E).

### Recurrent networks recapitulate the triplet motifs predicted by the three-neuron model

Our characterization of three-neuron circuits undergoing STDP revealed six distinct triplet motif classes: (1) no connections, (2) no bidirectional connections, (3) only unidirectional connections, (4) mixed unidirectional and bidirectional connections, (5) no unidirectional connections, and (6) only bidirectional connections. Together, these classes contained 14 of the 16 possible triplet motifs (Fig. 2E). Motifs combining absent, unidirectional, and bidirectional connections (Fig. 2E, gray bubble) could not be generated by any single STDP rule, regardless of the balance between LTD and LTP or their respective timescales.

We next asked whether these constraints persist when motifs are embedded in larger recurrent networks, where many pathways interact simultaneously (Fig. 3A). We simulated recurrent networks with excitatory and inhibitory neurons with plasticity at excitatory-to-excitatory synapses, and inhibitory plasticity controlling excitatory firing rates (Fig. 3B). In the presence of unstructured Poisson input, we quantified the resulting triplet motif distributions in the excitatory population. The steady-state distributions reproduced the same motif classes identified in the smaller three-neuron circuits (Fig. 3C). Across parameter choices, the simulations confirmed the analytical prediction that no single plasticity rule produces the full repertoire of triplet motifs (Fig. 3D).

**Figure 3:**
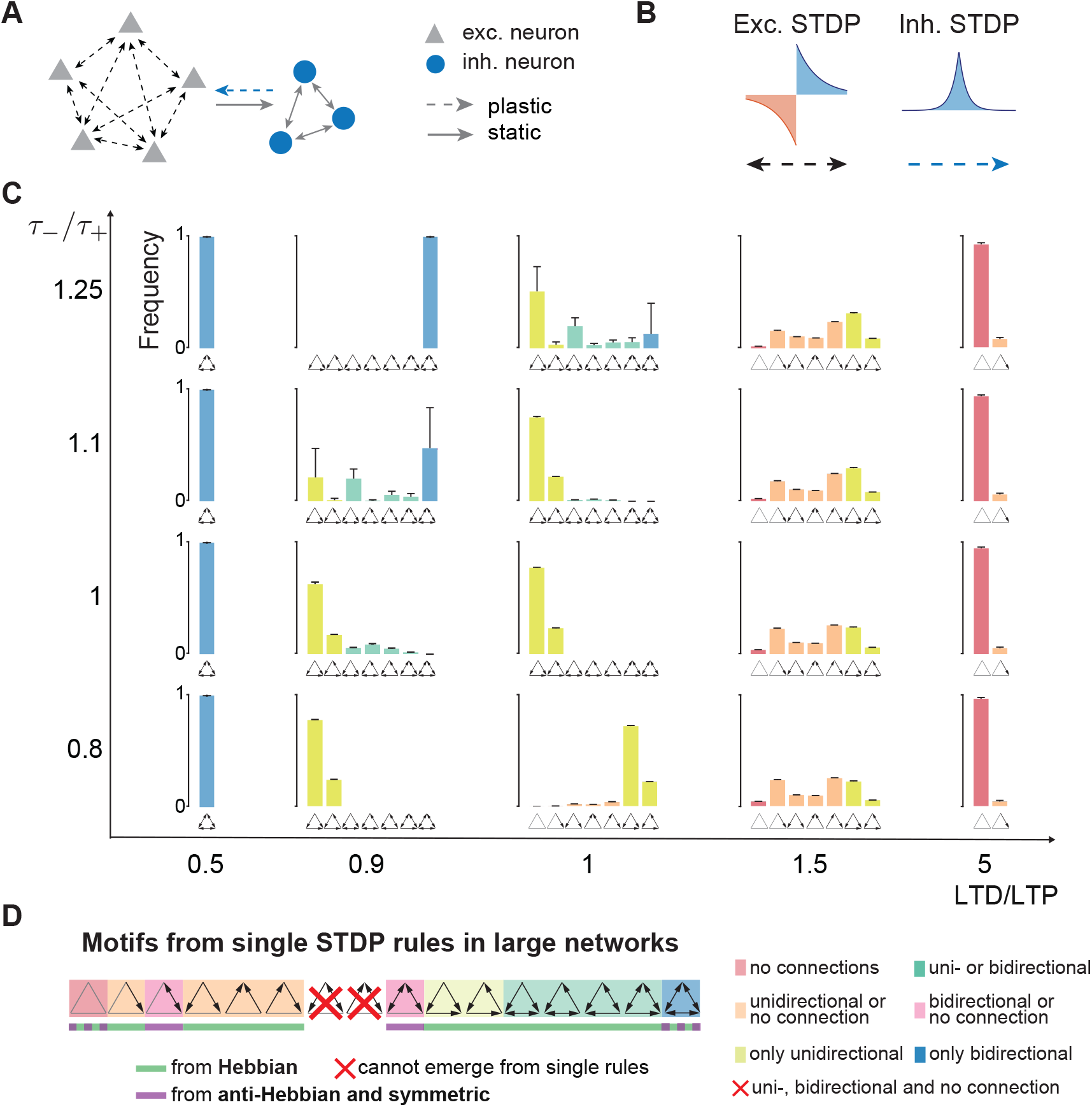
Recurrent networks recapitulate the triplet motifs predicted by the three-neuron model. **A.** Schematic of the large network simulations, including inhibitory neurons that homeostatically regulate excitatory firing rates with inhibitory STDP^51^. **B**. Plasticity rules implemented at E-to-E synapses (left) and I-to-E synapses (right). **C**. Steady-state triplet motif distributions in the excitatory population under Hebbian rules for different values of LTD/LTP and *τ*_−_/*τ*_+_. Bar colors correspond to the regions in Fig. 2. Bars show the mean and error bars the standard deviation (*n* = 10 trials). **D**. Summary of all possible motifs from single STDP rules as in Fig. 2E (see also Fig. S3).

The rule-to-motif mapping is therefore many-to-one: distinct plasticity rules can produce the same qualitative motif outcomes. Broad regions of the STDP parameter space map to the same motif class, despite substantial differences in the timing and relative strength of LTP and LTD (Fig. 2). Moreover, anti-Hebbian and symmetric rules produced largely overlapping classes of motifs across the parameter space. This degeneracy arises because motif formation depends not on the full STDP kernel itself, but on how the kernel determines the effective interaction coefficients in Eq. (1). Different STDP kernels can produce similar values, or the same sign patterns, of *M*_0_, *M*_10_, and *M*_01_, resulting in qualitatively similar trajectories in the synaptic weight phase plane and the same stable motifs.

Finally, motivated by experimental evidence that neuromodulation can alter the polarity and temporal profile of STDP rules^52–54^, we investigated how switching the plasticity rule affects motif distributions (Fig. S4, Fig. S5). After the network reached a steady-state motif distribution under one plasticity rule, we switched to a different rule. The new rule constrained the connectivity that emerged after the switch. Although the frequencies of individual motifs sometimes differed from simulations starting from random initial weights, the same motif classes emerged. Thus, the plasticity history of the network influences motif frequencies, while the currently active rule determines which motif classes can be maintained.

### Combining STDP rules gives rise to the full motif repertoire

Having shown that individual STDP rules impose strict constraints on the resulting triplet motifs, we next asked whether combining different rules could produce motifs absent under homogeneous plasticity. Biological circuits do not implement a single universal plasticity rule. Instead, distinct forms of plasticity coexist across synapse classes and neuron types^21^ and can change with modulatory state^52–54^. Therefore, we next hypothesized that the diversity of motif statistics may reflect the interaction of multiple plasticity rules operating within the same network. We then tested whether combining STDP rules could overcome the constraints imposed by any single rule and expand the repertoire of emerging triplet motifs.

To implement heterogeneity in the STDP rules, we assigned plasticity rules to synaptic connections based on the identity of the postsynaptic neuron (Fig. 4A). Under this configuration, all incoming synapses to a given neuron evolved under the same rule, while outgoing synapses from that neuron could follow distinct rules depending on the identity of their postsynaptic target. This choice was motivated by evidence that STDP expression is largely governed by postsynaptic mechanisms, such as calcium dynamics and backpropagating action potentials, and depends on the postsynaptic neuron type^12,21,55,56^. Once different synapses followed different learning rules, the effective synaptic change terms in Eq. (1) differed across connections, creating interactions that could not be reduced to any single homogeneous rule.

**Figure 4:**
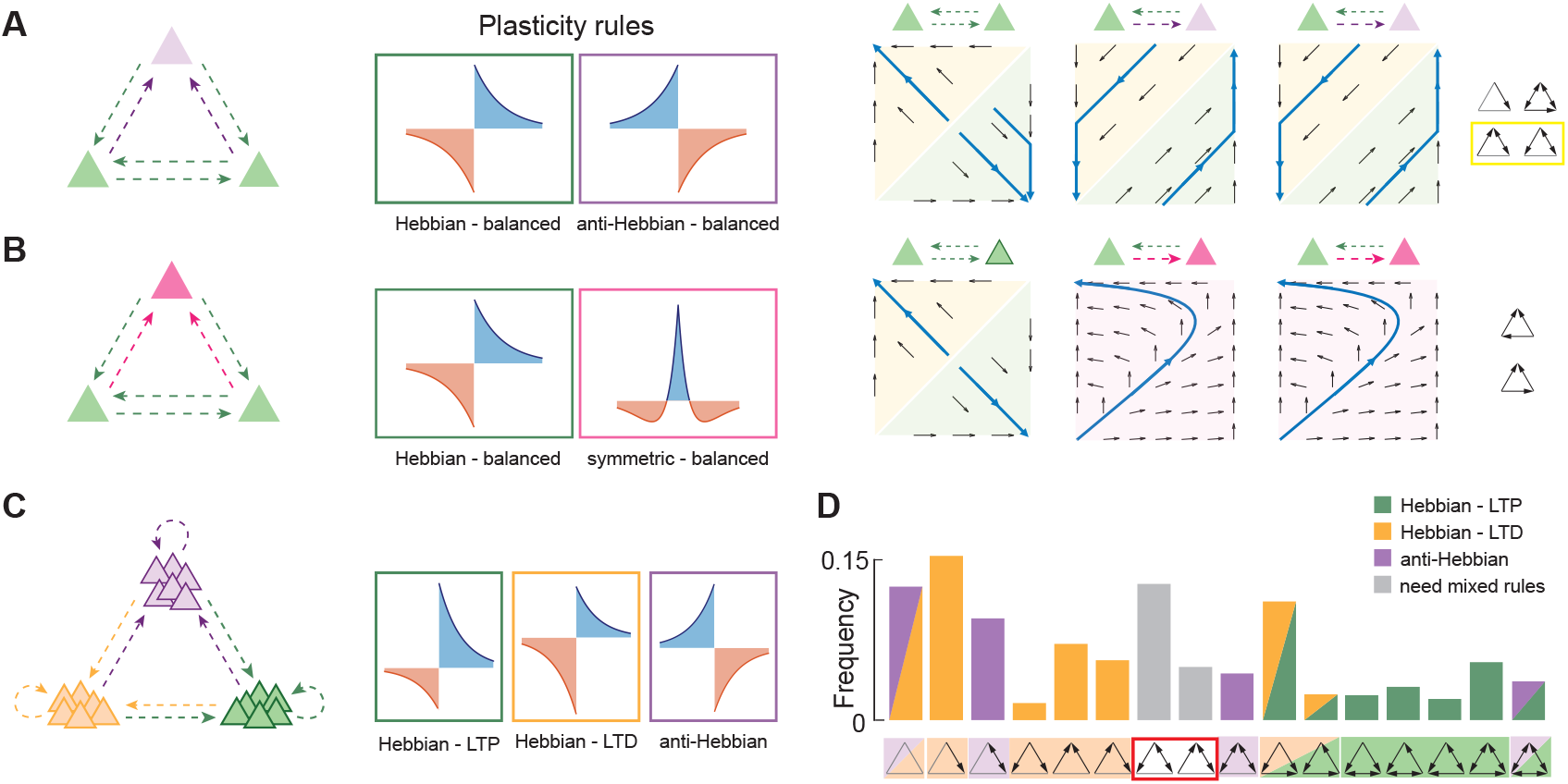
Heterogeneity in plasticity rules facilitates the formation of new motifs. **A**. *Left:* Connections between three neurons evolve according to Hebbian (green) or anti-Hebbian (purple) plasticity rules. *Right:* Phase planes for each pair of connections and the possible triplet motifs that emerge. Blue arrows show example trajectories. **B**. Same as in A, but for Hebbian (green) and symmetric (pink) STDP rules. **C**. Schematic of a large network with three plasticity rules: anti-Hebbian (purple), LTD-dominated Hebbian (orange), and LTP-dominated Hebbian (green). The network is initially all-to-all connected. **D**. Distribution of triplet motifs across the entire network. Single-color bars indicate motifs formed by three neurons belonging to the same excitatory subpopulation, and two-color bars indicate motifs formed spanning two excitatory subpopulations. Gray bars indicate motifs that require interactions between different plasticity rules.

As a minimal demonstration, we again considered a three-neuron circuit in which the incoming connections to two neurons followed a Hebbian rule and the incoming connections to the third neuron followed an anti-Hebbian rule. Both rules were chosen to have balanced LTD and LTP. When applied homogeneously to all connections in the circuit, each rule produced only a restricted set of motifs: for the chosen parameters, the Hebbian rule produced only unidirectional connections, while the anti-Hebbian rule produced bidirectional connections within the bounds (Fig. 2). When combined in the same circuit, the two rules produced new classes of motifs combining absent, unidirectional, and bidirectional connections, which could not be obtained from any individual STDP rule (Fig. 4A). Moreover, anti-Hebbian and symmetric rules, although producing the same classes of motifs when applied in isolation, led to different outcomes when combined with Hebbian plasticity. For example, combining a Hebbian rule with a symmetric rule led to a single unidirectional connection (Fig. 4B), but this was not the case when combining a Hebbian rule with an anti-Hebbian one.

Finally, we tested these predictions in large recurrent networks of excitatory and inhibitory neurons receiving unstructured Poisson inputs. We partitioned the excitatory population into three subpopulations of equal size, each evolving its incoming synaptic strengths according to a distinct STDP rule (Fig. 4C). In addition, the I-to-E synapses evolved under a homeostatic inhibitory plasticity rule that regulated excitatory firing rates^51^, allowing excitatory STDP to drive connectivity structure. Across the excitatory population, combining distinct STDP rules generated motifs beyond those formed within the individual subpopulations and produced the full repertoire of triplet motifs (Fig. 4D). Thus, the effects predicted from three-neuron circuits also emerged in larger recurrent networks.

### Heterogeneous STDP rules increase motif diversity but preserve degeneracy in the rule-to-motif mapping

Having established that heterogeneous STDP rules increase motif diversity, we next asked how this heterogeneity affects the rule-to-motif mapping compared with homogeneous rules. We therefore explored the space of heterogeneous STDP rules using Monte Carlo simulations of recurrent networks as before (Methods). We divided the excitatory population into three subpopulations of equal size and assigned an STDP rule family, Hebbian (*Hs*), anti-Hebbian (*A*), or symmetric (*S*), to the incoming connections of each subpopulation. This yielded ten possible rule combinations, denoted by the rule letters assigned to the three subgroups, for example *HHA* (Fig. 5A). For each combination, we ran 500 simulations, giving 5,000 simulations in total, and randomly sampled the STDP parameters of each rule (Methods). With the exception of *HAS*, which contains all three rule families, each combination was dominated by one family assigned to at least two of the three subpopulations. For example, *HHA* contained two Hebbian rules with different parameters, and therefore different shapes.

**Figure 5:**
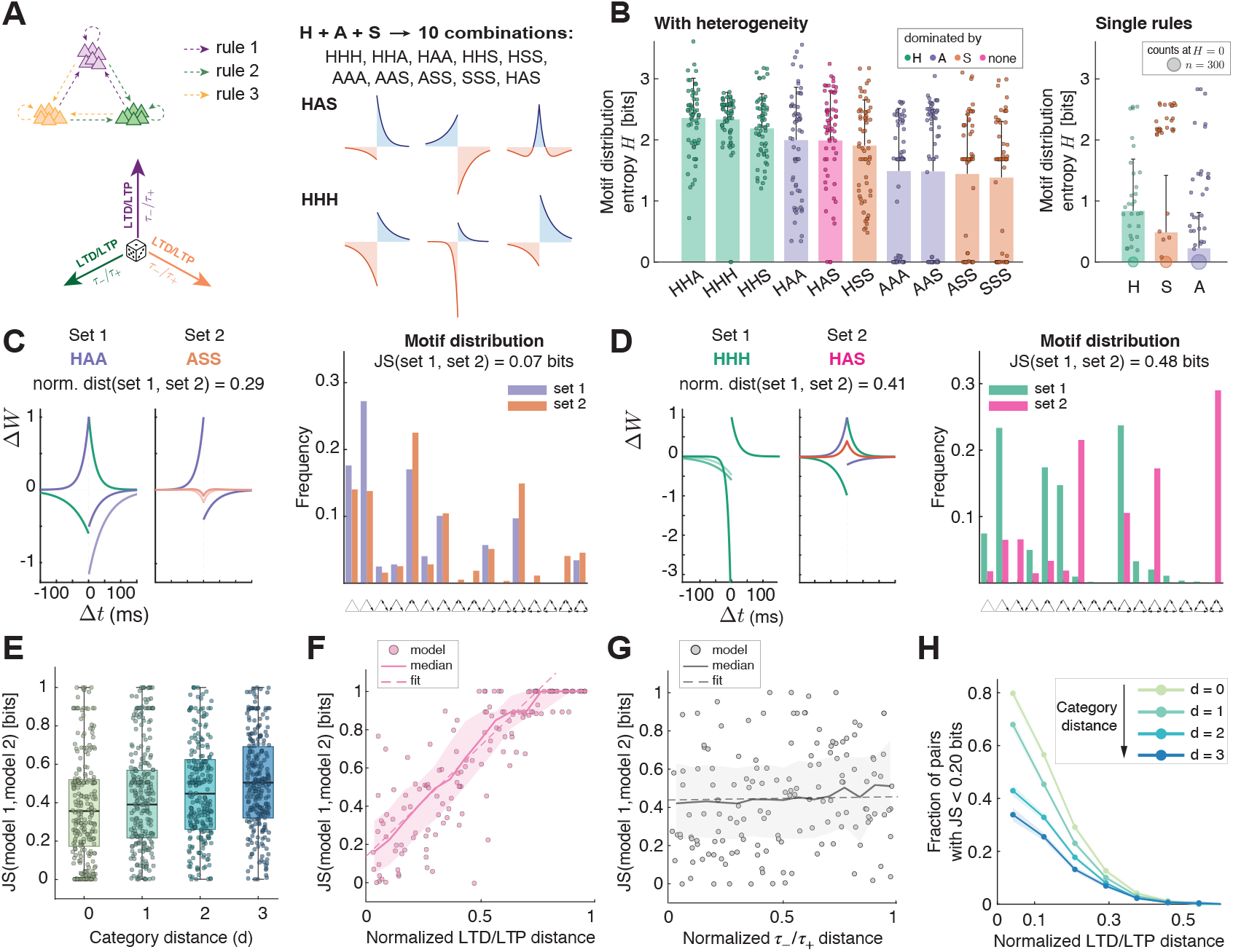
Heterogeneity increases motif diversity while preserving degeneracy in the rule-to-motif mapping. **A.** Schematic of the dataset generated from simulations with heterogeneous STDP rules. The excitatory population was divided into three subpopulations, each assigned an STDP rule with random parameters and deterministic family: Hebbian (*H*), anti-Hebbian (*A*), or symmetric (*S*). This yields ten possible categories (e.g., *HHA, AAS*), with 500 Monte Carlo simulations per category. **B**. Mean Shannon entropy of the steady-state motif distributions for each heterogeneous category (*left*) and for individual homogeneous rules (*right*). Bars show the mean, error bars the standard deviation, and points individual simulations. The area of the circle at entropy *H* = 0 is proportional to the number of simulations with zero entropy. Colors indicate the dominant rule family in each category. **C**. Example pair of simulations with low JS divergence (0.07 bits) despite a large distance between their rules (category distance *d* = 2) and parameters (normalized Wasserstein distance between log(LTD/LTP) of the three rule pairs, *W*_1_ = 0.29). **D**. As in C, but with higher JS divergence (0.48 bits) for a pair of simulations from different categories (category distance *d* = 2) and parameters (normalized Wasserstein distance between log(LTD/LTP) of the three rule pairs, *W*_1_ = 0.41). **E**. JS divergence between pairs of simulations as a function of category distance *d*, defined as the number of differing rule-family assignments. Boxes indicate the median and interquartile range (IQR), whiskers extend to 1.5 × IQR, and points show 250 example pairs. **F**. JS divergence as a function of the normalized Wasserstein distance between the log_2_(LTD/LTP) parameter values of each pair of simulations. Points show example pairs, solid line the moving median, shading the IQR, and dashed line the linear fit. **G**. Same as in **F**, but for the normalized Wasserstein distance between the log_2_(*τ*_−_/*τ*_+_) parameter values. **H**. Fraction of pairs with low JS divergence (<0.2 bits) as a function of the normalized Wasserstein distance between the log_2_(LTD/LTP) parameter values, separated by category distance *d*.

To quantify how heterogeneity affected the diversity of network structures after plasticity, we computed the Shannon entropy of the final motif distributions (Fig. 5B, left). All heterogeneous categories had higher mean entropy than the cases with a single STDP rule (Fig. 5B, right). Categories dominated by Hebbian rules produced the highest entropy, while categories containing only anti-Hebbian and symmetric rules produced the lowest. Nonetheless, even these categories had substantially higher entropy than the corresponding cases with a single rule, because different STDP parameters were assigned to the three subpopulations.

We further investigated how combining several rules affected the degeneracy of the rule-to-motif mapping. To dissect which factors contribute to degeneracy vs. specificity in the rule-to-motif mapping, we compared the simulations pairwise. We drew pairs of simulations from any of the ten categories and computed the Jensen-Shannon (JS) divergence^57^ (Methods): a symmetric dissimilarity measure bounded between 0 and 1, where 0 indicates identical distributions. We then related JS divergence to rule dissimilarity using three complementary distance measures: a category distance *d*, counting the number of rule families that differ between two simulations (e.g., *d* = 0 for simulations from the same category, such as *HHH* vs. *HHH, d* = 1 for *HHA* vs. *HHH*, etc.); a Wasserstein distance comparing the three LTD/LTP values; and a second Wasserstein distance comparing the three *τ*_−_/*τ*_+_ values (Methods). A low JS divergence at large rule distances indicates degeneracy because different rules converge to similar motif distributions (Fig. 5C). Conversely, an increase in JS divergence with rule distance indicates specificity because the motif distributions retain information about the underlying rules (Fig. 5D).

The JS divergence increased only weakly with category distance (Spearman’s correlation *ρ* = 0.17, permutation p < 0.001). Pairs from different categories were therefore not necessarily dissimilar, and pairs from the same category were not necessarily similar (Fig. 5E). Thus, the identity of rule families alone predicted motif dissimilarity only weakly. When analyzing specific parameter distances, we found that the LTD/LTP distance was the main predictor of motif dissimilarity (Fig. 5F; linear fit: *y* = 1.15 *x* + 0.15; Spearman’s correlation *ρ* = 0.59, permutation *p* < 0.001), while the *τ*_−_/*τ*_+_ distance was uncorrelated with the JS divergence across its entire range (Fig. 5G; linear fit: *y* = 0.02 *x* + 0.44; Spearman’s correlation *ρ* = 0.01, permutation *p* = 0.22). Accordingly, the fraction of pairs with low JS divergence (< 0.20) decreased as the LTD/LTP distance increased (Fig. 5H), indicating increasing specificity. However, even at an LTD/LTP distance of 0.3, a considerable fraction of model pairs retained a low JS divergence, particularly at smaller category distances (*d* = 0 or *d* = 1, between 20% and 30% of pairs; Fig. 5H). Thus, substantially different combinations of rules and parameters could still produce similar motif distributions, demonstrating degeneracy in the rule-to-motif mapping, in line with broader examples of degeneracy in neuronal and circuit models^58–60^.

In conclusion, degeneracy has two immediate consequences for interpreting motif distributions. First, observing a particular triplet motif distribution does not uniquely identify the underlying plasticity rules, but instead constrains them to a set of candidate mechanisms that produce similar motif statistics. Second, this nonuniqueness suggests a potential form of structural robustness, because similar local circuit structure can arise from different plasticity rules and parameter values.

### Inferring candidate plasticity rules from experimental motif distributions

We next considered the inverse problem: given a target motif distribution, can we identify which combinations of plasticity rules and parameters are compatible with it? Because the mapping from rules to motifs is degenerate, there is no unique answer; we therefore aimed to determine the range of combinations of rule families and parameter regimes that were compatible with a given motif profile. To illustrate this inverse mapping, we considered two target motif distributions obtained using different measurement modalities and at different scales: multipatch recordings from rat visual cortex^31^ and a connectomic reconstruction of mouse visual cortex^34^. We considered only motifs with connections involving all three neurons, excluding the unconnected motif and motifs involving only two-neuron interactions. For each target, we compared the simulated motif distributions described in the previous section with the experimental distribution using the Jensen-Shannon (JS) divergence.

Pooling across all STDP parameters, we analyzed how the JS divergence depends on the combination of STDP rule families (Fig. 6A, B left). For both datasets, combinations containing at least one Hebbian rule generally provided better fits, consistent with the prominent unidirectional structure generated by Hebbian plasticity. The fit also depended on the balance between LTD and LTP, quantified as the average of log(LTD/LTP) across the three subpopulations. For combinations with Hebbian rules, the JS divergence from the mouse dataset decreased steadily with this quantity and fell below 0.2 only for LTD-dominated parameters, whereas for combinations without Hebbian rules it remained close to 0.65 regardless of the LTD/LTP balance (Fig. 6A, right). For the rat dataset, JS divergence also decreased with the LTD/LTP balance, but remained above 0.3 in both groups and was lower for combinations with Hebbian rules at most values (Fig. 6B, right). However, the inverse mapping was not unique. Several distinct combinations of rule families produced comparably low JS divergence, and each combination covered a wide range of JS divergence values, reflecting substantial variability in the resulting motif distributions.

**Figure 6:**
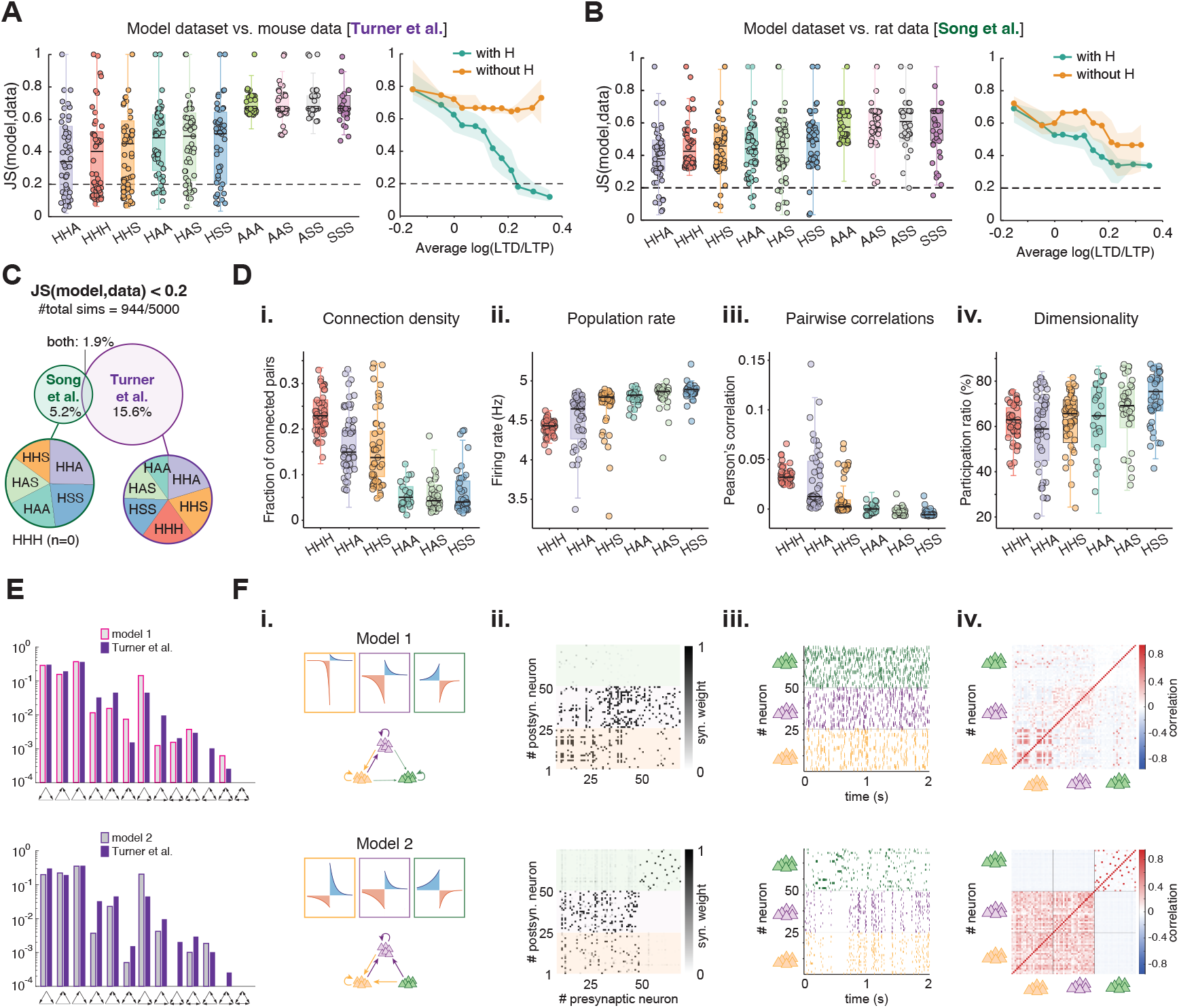
Heterogeneous STDP rules capture experimental motif distributions and shape network activity. **A.** JS divergence between model and mouse connectomic data^34^. Each network comprises three subpopulations, each following one STDP rule family among Hebbian (H), anti-Hebbian (A), symmetric (S); category labels denote the rule family of each subpopulation. *Left:* JS divergence for all ten categories, pooling across rule parameters. Categories are sorted by median JS divergence. Boxes show the median and interquartile range (IQR); whiskers extend to the most extreme values within 1.5 × IQR; points show a random subsample (10%) of individual networks. The dashed line marks the threshold for low JS divergence (0.2). *Right:* Median JS divergence (line) and IQR (shading) as a function of the average log_2_(LTD/LTP) across the three subpopulations, for categories with at least one Hebbian subpopulation (with H) and without (without H). Bins contain equal numbers of simulations. **B**. Same as in A, but for rat multipatch data^31^. Categories are ordered as in A. **C**. *Top:* Venn diagram of simulations with low JS divergence (< 0.2; 944 of 5000 in total) from the mouse data (15.6%, 780 simulations), the rat data (5.2%, 258), or both (1.9%, 94). *Bottom:* Composition of each subset by rule category. Categories without H are not shown: only 5 (mouse) and 37 (rat) of their simulations reached low JS divergence, compared with 775 and 221 simulations from categories with H, respectively. No HHH simulation matched the rat data. **D**. Network and activity properties across rule categories with at least one H subpopulation, for simulations with JS divergence from the mouse data below 0.2: **i)** connection density, **ii)** population firing rate, **iii)** mean pairwise Pearson correlation, and **iv)** dimensionality, quantified as participation ratio. Boxes show the median and IQR; whiskers extend to the most extreme values within 1.5 × IQR; points show a random subsample (30%) of individual networks. **E**. Frequency of three-neuron connectivity motifs (log scale) for two exemplar networks from our dataset (model 1 and model 2), each closely matching the mouse motif distribution^34^. **F**. Structure and activity of the simulated networks in E. *Top:* model 1, with rules: Hebbian with *θ* = 2.83, *γ* = 0.40; Hebbian with *θ* = 2.77, *γ* = 1.49; anti-Hebbian with *θ* = 1.17, *γ* = 1.53. *Bottom:* model 2, with rules: Hebbian with *θ* = 2.71, *γ* = 2.95; Hebbian with *θ* = 2.05, *γ* = 0.76; anti-Hebbian with *θ* = 2.67, *γ* = 2.36. **i**. STDP rules and population graphs. **ii**. Connectivity matrices after convergence of synaptic weights. Background colors refer to the rule assigned to each subpopulation. **iii**. Spiking activity of the excitatory subpopulations after convergence; colors as in the weight matrix. **iv**. Pearson correlation of binned spike counts across all neuron pairs.

Focusing on simulations with low JS divergence, JS(model,data) < 0.2, we found that almost 20% of all simulations (944/5000) produced a motif distribution similar to at least one of the two datasets. Specifically, 15.6% of all simulations were similar to the mouse dataset^34^, 5.2% to the rat dataset^31^, and 1.9% to both (Fig. 6C). Categories without Hebbian rules almost never reached this threshold (5 simulations for the mouse dataset and 37 for the rat dataset, out of about 2000 simulations without Hebbian rules), while all categories containing at least one Hebbian rule were represented at similar frequencies. However, while the category containing only Hebbian rules (*HHH*) accounted for a large fraction of the good fits to the mouse dataset, it did not fit the rat dataset (Fig. 6C). Thus, low-divergence fits strongly favor combinations containing at least one Hebbian rule, while the best-performing rule composition depends on the dataset.

To understand what drove these low-divergence fits, we next asked whether they reflected detailed agreement in motif frequencies or more general properties of the connectivity, particularly its density. Both target distributions were dominated by motifs with relatively few connections. Consequently, some simulations achieved a low JS divergence primarily by producing sparse connectivity rather than by closely matching the frequencies of the less common motifs (Fig. 6Di). A low JS divergence therefore does not identify a unique network structure.

In sum, these observations indicate degeneracy in the rule-to-motif mapping, because multiple combinations of plasticity rules can produce similar motif statistics. Therefore, rather than identifying a single best rule, our approach reveals a set of plausible plasticity mechanisms consistent with the observed motif distribution.

Next, we investigated how the categories defined by different combinations of plasticity rules affected activity in the resulting networks. We measured the population firing rate, mean pairwise correlations, and dimensionality after plasticity. We restricted our analysis to simulations with low JS divergence from the mouse data (Fig. 6D), which all contained at least one Hebbian rule. Population firing rates varied little in absolute terms across categories and remained close to, but below, the homeostatic target of 5 Hz, consistent with the homeostatic plasticity rule stabilizing excitatory activity around this value. Mean pairwise correlations were low for most rule categories but increased in the category containing only Hebbian rules (*HHH*), which differed significantly from all other categories except *HHA* (Dunn–Šidák post-hoc, all *p* < 10^−13^ ). The *HHA* and *HHS* rule categories additionally generated broad distributions of mean pairwise correlations, with several highly correlated networks. Dimensionality, defined by the participation ratio^61^ (Methods), was high across all categories, consistent with the unstructured Poisson input and the sparse connectivity of networks matching the experimental data, in which many triplets contained absent connections. Differences across categories were statistically significant for all three measures (Kruskal–Wallis test, all *p* < 10^−17^). The rule category accounted for a large fraction of the rank variance in population firing rate (*ϵ*^2^ = 0.59) and pairwise correlations (*ϵ*^2^ = 0.67), but only for a small fraction in dimensionality (*ϵ*^2^ = 0.12). Thus, networks with similarly good matches to the motif distribution can nevertheless show different firing rates and pairwise correlations depending on the combination of plasticity rules, whereas dimensionality varies much less.

To illustrate the variability within a single rule category, we selected two *HHA* networks from the broad range of activity patterns described above, both of which closely matched the mouse motif distribution (Fig. 6E). Both networks combined two Hebbian and one anti-Hebbian subpopulation, but with different rule parameters. Despite the similarity in their motif distributions, the two networks differed in their activity and in the density and organization of connections within and across subpopulations (Fig. 6F). The first network showed asynchronous firing in all three subpopulations, and pairwise correlations were weak apart from a block of positively correlated neurons in one Hebbian subpopulation. In contrast, the second network showed sparser and bursty activity, and pairwise correlations were strongly positive across the two Hebbian subpopulations, whereas the anti-Hebbian subpopulation was only weakly correlated with them. These examples demonstrate that networks with nearly identical motif statistics can operate in markedly different dynamical regimes.

Together, these results show that experimentally observed motif distributions are compatible with several combinations of plasticity rules. Although these combinations are degenerate at the level of motif structure, they can differ in their firing rates and pairwise correlations.

## Discussion

In this work, we established an analytical rule-to-motif map linking excitatory spike-timing-dependent plasticity (STDP) rules to the emergence of local recurrent connectivity. By deriving motif-specific interaction coefficients from the plasticity rule and spike-train covariances, we obtained a mapping between the parameter space of excitatory plasticity rules and the resulting steady-state triplet motifs. Individual Hebbian, anti-Hebbian, and symmetric rule families imposed strong restrictions on the connectivity patterns that could emerge. In contrast, combining different excitatory rules allowed all 16 triplet motifs to emerge in larger recurrent networks. The resulting mapping from rules to motifs was strongly degenerate, with distinct rules and combinations of rules producing similar motif distributions. Thus, circuit structure carries information about the underlying plasticity mechanisms, but generally narrows the possibilities rather than identifying a unique rule.

Previous theoretical work has established that the shape of excitatory STDP rules can strongly influence the structure that emerges in recurrent networks. Hebbian rules have been studied extensively in the context of formation and stability of recurrent connectivity^19,20,62,63^. More recently, the degree of temporal causality of STDP has been varied to show how rule shape controls the stability of overlapping neuronal assemblies^27^. Previous studies often started from prespecified STDP rules and asked which structures they generate^45,49,64,65^. Here, we systematically explored how different families of STDP rules and their parameters constrain the repertoire of local connectivity motifs that can emerge. Within each family of STDP rules, the balance between LTD and LTP mainly determined the class of motifs, although their relative time constants shifted the boundaries between regimes and could change the resulting motifs. For example, when total LTD and LTP were balanced, bidirectional connections emerged under a Hebbian rule if the LTD time constant was sufficiently larger than the LTP time constant. Consistent with this, when different STDP rules were combined, differences in LTD/LTP balance predicted differences in the distributions of motifs much better than differences in either the combination of families of rules or the relative LTD and LTP time constants. This comparison relies on relatively simple measures of distance between rules, and alternative measures that capture additional features of the STDP rules may reveal further relationships. Beyond these effects of the parameters, combining different STDP rules produced outcomes that could not be predicted from the behavior of individual rules. For example, anti-Hebbian and symmetric rules produced largely overlapping classes of motifs in networks governed by a single family of rules, yet generated different outcomes when combined with Hebbian plasticity. Thus, combining different STDP rules does not simply produce intermediate outcomes between those generated by individual rules, but can give rise to connectivity patterns that do not emerge when the rules act individually.

Our results suggest that specific motifs do not uniquely identify the plasticity rule that generated them, but can instead emerge from multiple rules. Similar many-to-one relationships have been described for intrinsic conductances, where distinct parameter combinations can generate similar neuronal activity^66^, and more broadly across neuronal and circuit models^58–60^. This degeneracy became more pronounced when considering heterogeneous combinations of rules and parameters, where substantially different rules and parameter values could produce very similar motif distributions. Including such heterogeneity in plasticity rules supported a wide variety of motif distributions, including connectivity statistics measured in distinct experiments^31,34^. This degeneracy was nonetheless constrained in two ways. First, Hebbian plasticity provided an important constraint on matching the experimental motif distributions, as combinations without Hebbian rules rarely achieved low JS divergence for either dataset, consistent with the prominent unidirectional structure generated by Hebbian plasticity. Second, among combinations containing Hebbian rules, low JS divergence from the mouse dataset was reached predominantly for LTD-dominated parameters, whereas balanced or LTP-dominated parameters gave poorer fits. In both experimental datasets, sparse motifs containing few connections were much more frequent than densely connected motifs. As a result, simulations could match the experimental distributions mainly by reproducing this overall sparsity, even when they reproduced the frequencies of rarer motifs less accurately. The experimental comparison therefore constrains the set of candidate plasticity mechanisms that are compatible with the observed structure, but does not uniquely identify which rules generated these circuits. Such degeneracy could contribute to structural robustness by allowing similar connectivity patterns to arise from different plasticity mechanisms.

Degeneracy at the level of motif structure did not imply identical network dynamics. Among simulations that closely matched the mouse motif distribution^34^, different sets of Hebbian, anti-Hebbian, and symmetric STDP rules generated similar local connectivity motifs but produced systematic differences in population firing rate and pairwise correlations. Thus, different sets of STDP rules can produce distinct activity patterns despite very similar motif distributions. This suggests that connectivity and activity provide complementary constraints on the underlying plasticity mechanisms. Activity measurements do not uniquely identify the rules in our model, but they can further constrain the candidate rules that remain compatible with the circuit structure. More generally, population activity and perturbation responses can carry information about underlying plasticity mechanisms and recurrent network organization^67,68^.

The scope of our work was to provide a mechanistic understanding of how diverse plasticity rules shape self-organization in recurrent circuits. To achieve this, we adopted specific modeling choices that balance biological plausibility with analytical tractability. Networks started from random all-to-all connectivity and received homogeneous external input, allowing us to isolate connectivity structure generated by plasticity rather than by pre-existing topology or externally imposed correlations. In contrast to models in which assemblies are imposed by selectively increasing firing rates or correlations within predefined groups^69–71^, the external input in our model did not prescribe which neurons should become preferentially connected. This allowed connectivity structure to emerge spontaneously through plasticity, as in related work on autonomous self-organization^45,72^. An important extension will be to determine how constraints already present in the initial connectivity, such as the decrease in connection probability with intersomatic distance, interact with STDP to shape the resulting motifs^32,73,74^. We also used neuronal dynamics described by Hawkes processes and hard synaptic bounds. Additional mechanisms, such as synaptic normalization or intrinsic plasticity, could alter the long-term evolution of connectivity^45,63,69,75,76^. Although the Hawkes description provides analytical tractability, it omits nonlinear spike generation and refractory periods, which could alter the mapping from STDP rules to motifs in networks of leaky integrate-and-fire neurons.

Heterogeneity was implemented using three excitatory subpopulations, with incoming synapses assigned according to postsynaptic neuron identity, motivated by evidence that STDP expression often depends on the postsynaptic neuron type^12,21^. Alternative assignment schemes are likely to generate different motif statistics. In addition, our analysis focused on excitatory-to-excitatory triplet motifs while inhibitory plasticity provided homeostatic regulation of excitatory activity^51^. Experimental studies have revealed highly structured patterns of connectivity among inhibitory neurons and between excitatory and inhibitory neurons^77–79^. Diverse inhibitory plasticity rules can themselves generate such structured connectivity^72,80–82^, suggesting that interactions between excitatory and inhibitory plasticity will further expand the space considered here. Finally, experimental studies show that neuromodulators can alter the polarity and temporal window of STDP^52,53^, and that these effects can depend on recent activity at individual synapses^54^. Here, we model neuromodulation as a global change in the plasticity rule.

Although we modeled three excitatory subpopulations, the same analysis can be extended to larger numbers of rule combinations. A more extensive exploration of heterogeneous combinations of rules could be combined with simulation-based inference or meta-learning approaches to identify sets of plasticity mechanisms compatible with increasingly detailed experimental constraints^83–85^. Beyond structural and activity statistics, such approaches could also constrain plasticity rules by the computations or behavioral signals they support, such as familiarity or novelty detection, contextual novelty, or memory-related dynamics^86^. This would allow candidate plasticity mechanisms to be tested not only against circuit structure and spontaneous activity, but also against network function.

In conclusion, our analysis links local STDP rules to the emergence of connectivity motifs. Rather than pointing to a single plasticity mechanism underlying an observed circuit structure, our results show that distinct rules can generate similar motif distributions while differing in their firing rates and pairwise correlations. Combining structural and functional measurements may therefore provide a way to more tightly constrain the plasticity mechanisms consistent with biological circuit data.

## Methods

To study triplet connectivity motifs arising from different STDP rules, we considered three excitatory neurons connected with plastic synapses. In this section, we introduce neuronal dynamics and our analysis for the evolution of three-neuron connectivity structures.

### Neuronal dynamics

We considered networks of *N* excitatory neurons, recurrently connected through synaptic weights defined in the connectivity matrix *W*, without self-connectivity (i.e. *w*_*ii*_ = 0), where rows (columns) denote post-(pre-) synaptic neuron indices, so the connection from a presynaptic neuron *j* to a postsynaptic neuron *i* is *w*_*ij*_. The spike train *S*_*i*_(*t*) of a neuron *i* at time *t* can be written as a sum of Dirac deltas:

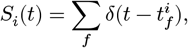

where 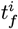 denotes the times that neuron *i* has spiked. We modeled the activity of a neuron *i* as a Hawkes process, namely, an inhomogeneous Poisson process whose intensity (i.e. expected firing rate) is given by:

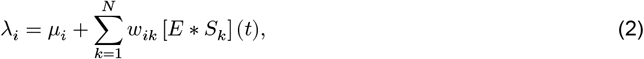

where *μ*_*i*_ represents a constant external input firing rate and *E*(*t*) is the excitatory postsynaptic potential (EPSP), which we assumed to be identical for all neurons:

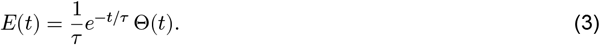

The symbol * denotes the convolution operator:

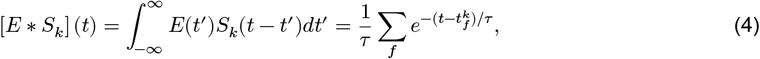

where the last equation follows from the properties of the Dirac delta.

### Synaptic weight dynamics

The framework used in this section to describe synaptic weight dynamics under STDP follows previous work on plasticity in recurrent spiking networks^16,45,47,49^. We summarize it here for completeness and to define notation.

The synaptic evolution of a three-neuron structure corresponds to a dynamical system of six equations (two equations for each pair of neurons, seen both as pre- and postsynaptic). For each connection *w*_*ij*_, the synaptic change 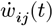 due to a spike timing-dependent plasticity (STDP) rule can be written as:

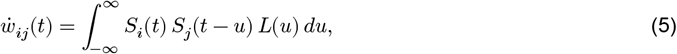

where *S*_*j*_(*t* − *u*) and *S*_*i*_(*t*) are the spike trains of the pre- and the postsynaptic neuron, respectively, and *L*(*u*) denotes the STDP function. We further impose hard bounds

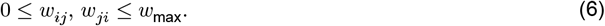

to prevent unbounded growth or negative weight. Assuming separation of timescales between (slow) synaptic plasticity and (fast) neuronal dynamics^16^, the time-averaged synaptic change 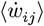 can be written as the integral of the product between the correlation function of the pre- and postsynaptic spike trains and the STDP function:

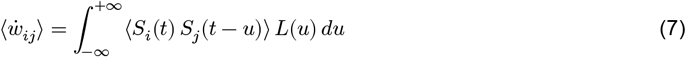

Or, equivalently,

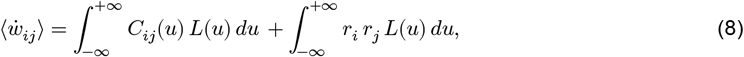

where *r*_*i*_ and *r*_*j*_ are the mean firing rates of neuron *i* and neuron *j*, and *C*_*ij*_ is their covariance density. Note that others may refer to the quantity *C*_*ij*_ as the correlation function. Here, the correlation is defined as the sum *r*_*i*_ *r*_*j*_ + *C*_*ij*_(*u*). Intuitively, the correlation represents the probability that neurons *j* and *i* fire with a delay in time *u*. Previous work has extensively described how the Fourier transform of the covariance *C*_*ij*_ can be expressed as a power series that depends on the full network connectivity *W* and neural activity *r*_*k*_^47,49^:

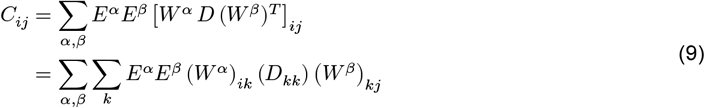

where *D* is a diagonal matrix with the mean firing rates *r*_*k*_ of all neurons on its diagonal, and *α, β* represent the number of synaptic connections in the path from neuron *k* to *i* and from *k* to *j*.

Following ref. ^45^, as a consequence of Eq. (8), we can expand the average change in synaptic weight into the sum of terms that depend on connectivity, activity, and plasticity. More specifically, Eq. (8) can be re-written as

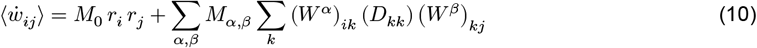

where *M*_0_, *M*_*α,β*_ are called “motif coefficients”, and correspond to *M*_0_, *M*_10_and *M*_01_in Eq. (1). These coefficients combine the STDP function with the neuronal activity of *i* and *j* triggered by spikes from a source neuron *k*, and are indexed by the number *α, β* of synaptic connections from the source neuron to neuron *i* and *j*, respectively. The sum *α* + *β* is called the order of the motif coefficient. For *α* + *β* = 0, the term *M*_0_ refers to the contribution to the synaptic change given by the mean firing rates of neuron *i* and *j*, and is simply given by the integral of the STDP function:

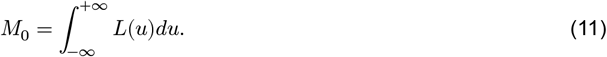

For *α* + *β* ≥ 1, the terms *M*_*α,β*_ are defined in the Fourier domain as

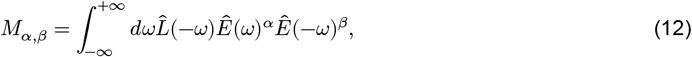

where 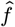 represents the Fourier transform of a function *f* . These terms refer to the contribution to the change in *w*_*ij*_ given by spikes traveling *α* (*β*) connections throughout the network from a source neuron *k* to the postsynaptic (presynaptic) neuron *i* (*j*). Therefore, besides the STDP function, they also include powers of the EPSP function. In this work, we consider the power series in Eq. (10) truncated to the first order, namely, we take into account only spikes from the neurons involved in the given connection *w*_*ij*_:

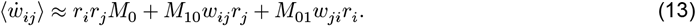

The terms *M*_10_ and *M*_01_ represent the contribution to the change 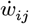 due to presynaptic and postsynaptic spikes, respectively. Since this approximation ignores terms of higher orders, we can break down the synaptic evolution of a three-neuron system into three dynamical systems, each for one pair of connections:

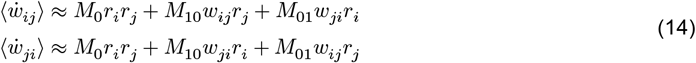

The coefficients *M*_0_, *M*_10_ and *M*_01_ depend on the specific shape of the STDP function *L*.

### Steady-state motifs under different STDP rules

We assumed constant firing rates, and analyzed the influence of the *M* coefficients on Eq. (14), depending on the STDP function. We analyzed the evolution of three dynamical systems for three types of STDP functions: Hebbian, anti-Hebbian, and symmetric rules. We calculated the fixed points, or attractors, for the system above given the bounds Eq. (6). The attractors are given by the intersection points of the nullclines and by upper (lower) bounds with non-negative (non-positive) derivative. There are, in principle, three possible types of attractors: no connection, unidirectional connection, and bidirectional connection. While for Hebbian rules attractors lie at the bounds (weights at either 0 or *w*_max_), for anti-Hebbian and symmetric rules, bidirectional attractors can also lie within the bounds (weights below *w*_max_), and for symmetric rules they can form a line of fixed points whose endpoints, where one weight reaches the lower bound, are unidirectional connections. If the STDP rule is the same for all pairs of connections, the resulting triplet motif is the combination of attractors from three identical phase planes. Therefore, for example, if the phase plane for a specific STDP rule shows bidirectional connections as the only possible attractor, then the only possible steady state motif will be all-to-all connected. If, instead, the STDP rule leads to either bi- or unidirectional connections, then there will be 7 possible motifs at the steady state, given by all possible combinations of uni- and bidirectional connections. The case of different STDP rules will be covered later.

### Asymmetric STDP rules

The asymmetric Hebbian STDP rule is defined as:

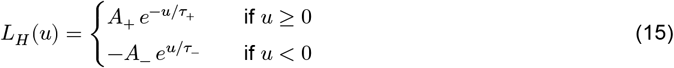

The asymmetric anti-Hebbian STDP rule is defined as:

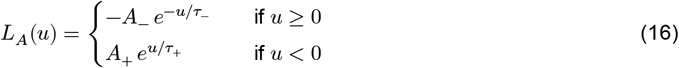

The parameters *A*_+_ > 0 and *A*_−_ > 0 denote the maximum amplitudes of LTP and LTD, respectively, and *τ*_+_ and *τ*_−_ their corresponding decay time constants.

For the Hebbian STDP rule, the coefficients *M*_0_, *M*_10_, *M*_01_ in Eq. (14) are the following (see refs. ^45,72^):

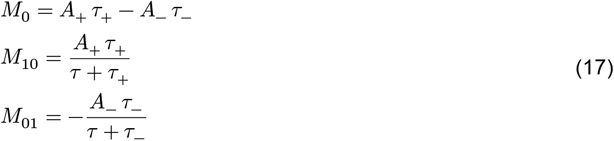

where *τ* is the EPSP time constant.

To simplify the notation in Eq. (14), we defined *A* := *A*_+_*τ*_+_. Moreover, we considered the ratio between the time constants *γ* = *τ*_−_/*τ*_+_, and the ratio between the overall amount of LTD and LTP, *θ* = *A*_−_ *τ*_−_/*A*_+_ *τ*_+_. Therefore, we can rewrite the coefficients for the Hebbian rule as:

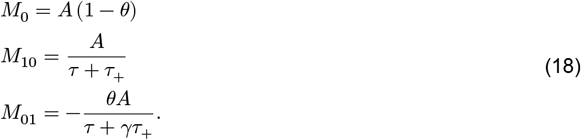

The resulting system is:

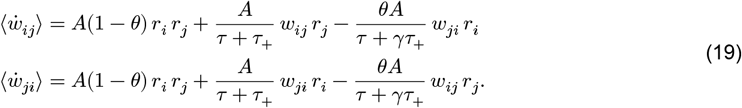

with 0 ≤ *w*_*ij*_, *w*_*ji*_ ≤ *w*_max_.

For the anti-Hebbian rule, the same parametrization applies, with *M*_10_ and *M*_01_ exchanged, while *M*_0_ remains unchanged.

### Symmetric STDP rule

The symmetric STDP rule was defined as:

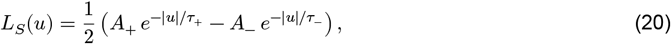

with *A*_+_, *A*_−_ > 0.

The coefficients for this rule are (cf.^72^)

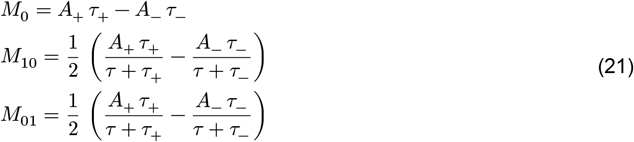

By defining as before *A* := *A*_+_*τ*_+_, *γ* = *τ*_−_/*τ*_+_, and *θ* = *A*_−_ *τ*_−_/*A*_+_ *τ*_+_, we can rewrite Eq. (21) as

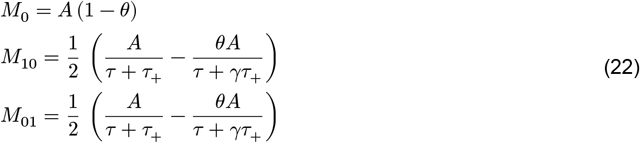

and

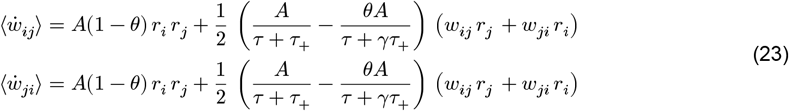

with

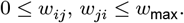

Eqs. (19) and Eqs. (23) allowed us to find regions in the parameter space (*θ, γ*) defined by the type of possible attractors. The boundaries for the regions in the parameter space depend on several factors besides the STDP parameters, including the upper bound on the weights and the firing rates (Supplementary Information, Section S1). Unless otherwise specified, the phase-plane analyses and analytical boundaries shown in Fig. 2 (and derived in Supplementary Information, Section S1) use the following baseline parameters: *w*_max_ = 0.5, *r* = 15 Hz, *τ* = 10 ms, *τ*_+_ = 20 ms. In particular, the firing rates may vary with time, and so the phase planes will also vary through the dynamics. However, the triplet motifs at the steady state are not significantly affected: even if the nullclines might change, the types of attractors either stay the same, or they become a subset of the initially predicted attractors. For simplicity, we assumed *r*_*i*_ = *r*_*j*_ =: *r* (corresponding to *C* = 1 in Supplementary Information). This assumption is approximately satisfied in the numerical simulations, since the homeostatic inhibitory plasticity rule we employed^51^ (see below) drives excitatory firing rates toward a common target rate.

### Triplet motif simulations under single STDP rules

To validate the analytical predictions in Fig. 2A, C, D, we simulated three-neuron circuits with Hawkes-process dynamics (Eq. (2)). Each of the three excitatory neurons was paired one-to-one with a dedicated inhibitory neuron: the excitatory-to-inhibitory connection was fixed and non-plastic (*w*_ie_ = 2.0), while the inhibitory-to-excitatory connection evolved under the homeostatic inhibitory plasticity rule^51^ with *τ*_vogels_ = 40 ms, target postsynaptic rate 15 Hz, initialized at *w*_ei_ = 5.0. No inhibitory-to-inhibitory connections were present. Excitatory-to-excitatory synapses evolved under the STDP rule being tested and were initialized uniformly in 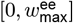, with 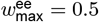. All neurons received independent, unstructured Poisson external input, at a fixed rate of 20 Hz for excitatory and 50 Hz for inhibitory neurons. The excitatory and inhibitory postsynaptic-potential time constants were *τ*_*e*_ = *τ*_*i*_ = 10 ms, and the STDP time constant *τ*_+_ = 20 ms.

For the Hebbian rule (Fig. 2B), we sampled *θ* ∈ [0.1, 2.5] and *γ* ∈ [0.4, 2.0], both with discretization step 0.1. For the anti-Hebbian and symmetric rules (Fig. S3), we sampled *θ* ∈ [0.8, 1.2] (step 0.01) and *γ* ∈ [0.8, 1.2] (step 0.05). For each combination of *θ* and *γ*, we ran *n* = 10 trials. Each excitatory-to-excitatory connection was classified as present at steady state if *w*_*ij*_ > 0.12 and absent otherwise, and the resulting triplet motif for each trial was assigned to one of the sixteen classes. Motif frequencies were averaged across trials for each parameter combination.

### Heterogeneity in the plasticity rules in triplets

To implement heterogeneity in the plasticity rules (Fig. 4), we assigned different types of rules based on the postsynaptic neuron. In this case, the synaptic evolution could vary across different pairs of connections. For example, we combined Hebbian and anti-Hebbian plasticity rules for three neurons *i, j, k*. Neuron *j* had incoming connections with anti-Hebbian plasticity, and neurons *i* and *k* had incoming connections with Hebbian plasticity. We chose both rules balanced in LTD/LTP (*θ* = 1, *γ* = 1). The synaptic evolution for *w*_*ik*_ and *w*_*ki*_ followed the same rule (Hebbian in both directions), but for *w*_*ij*_ and *w*_*ji*_ (and analogously for *w*_*jk*_ and *w*_*kj*_):

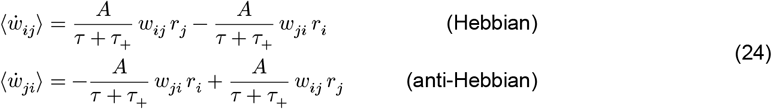

The system above has two attractor points, (*w*_max_, *w*_max_) and (0, 0). When we combined the possible attractors with the ones given by the Hebbian-rule system, we obtained the new motifs as in Fig. 4A.

Similarly, when we combined the same Hebbian rule with a symmetric rule with *θ* = 1, *γ* = 2, we obtained:

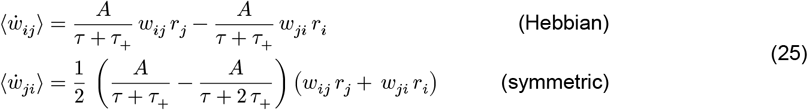

which led to a single attractor (0, *w*_max_), and the motifs as in Fig. 4B.

### Large network simulations

To study the emergence of triplet motifs on a larger scale, we considered neural networks of *N* = 100 neurons, of which *N*_*E*_ = 75 were excitatory (E) and *N*_*I*_ = 25 were inhibitory (I). All neurons received external Poisson input spike trains at fixed rate of 10 Hz.

The initial network was all-to-all connected (with no autapses). E-to-E synaptic weights were initially drawn from the uniform distribution over the interval 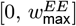, with 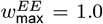, and followed excitatory STDP rules as described above, with time constant *τ*_+_ = 20 ms. I-to-E synaptic weights were initialized at *w*_ei_ = 5.0 and evolved to ensure homeostasis in excitatory dynamics^51^, with time constant *τ*_vogels_ = 40 ms and a target rate of 5 Hz. I-to-I and E-to-I connections were non-plastic (*w*_ii_ = 1.0, *w*_ie_ = 2.0). The excitatory and inhibitory postsynaptic-potential time constants were *τ*_*e*_ = *τ*_*i*_ = 10 ms. Simulations were run until the stopping criterion was reached, corresponding to a total of 10^8^ spikes emitted by the population, to ensure convergence of the synaptic weights. For the simulations shown in Fig. S4 and Fig. S5, the stopping criterion was instead set to 1.25 × 10^8^ spikes, corresponding to approximately 250,000 s, with the STDP rule switched after 110,000 s (about 30 hours).

To study the motif distribution after plasticity, we considered the connectivity submatrix of E-to-E connections. Each connection was classified as present at steady state if *w*_*ij*_ > 0.12 and absent otherwise. We calculated the frequency of each steady state motif based on previous approach^87^.

### Heterogeneity in the plasticity rules in larger networks

To implement heterogeneity in larger networks as in Fig. 4–Fig. 6, we divided the excitatory population into three subpopulations of equal size (25 neurons each), and assigned STDP rules according to the postsynaptic neuron, as in the three-neuron system. Thus, all incoming E-to-E synapses to neurons in the same subpopulation followed the same rule. In Fig. 4 C-D, we implemented an anti-Hebbian rule and two different Hebbian rules (parameters in Table S1).

To create our dataset in Fig. 5 and Fig. 6, we collected 5,000 Monte Carlo simulations, divided into 10 categories, each defined by a combination of Hebbian (*H*), anti-Hebbian (*A*) and symmetric (*S*) rules: *AAA, AAS, ASS, HAA, HAS, HHA, HHH, HHS, HSS, SSS*. Each category contained 500 simulations, with independently sampled STDP parameters (*θ*_*i*_, *γ*_*i*_)_*i*=1:3_ for the three rules. For each simulation, *θ* and *γ* for each assigned rule were sampled independently from the uniform distribution on [0.25, 3.0].

To quantify the diversity of the motif distribution, we computed its Shannon entropy Fig. 5B^88^:

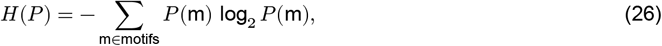

where [*P* (m_1_), …, *P* (m_16_)] is the vector containing the motif frequencies, measured after thresholding the weights (*w*_*ij*_ > 0.12).

To quantify the similarity between two motif distributions, we computed the Jensen-Shannon (JS) divergence (Fig. 5 and Fig. 6,^57^):

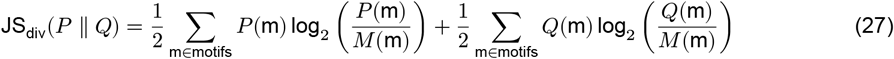

where *P* (m) and *Q*(m) are the two distributions, and 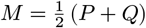.

To quantify the dissimilarity in rule-family composition between two simulations, we defined a category distance *d* as the number of rule families (out of three) that differ between the two simulations (e.g., *d* = 0 for two simulations from the same category, and *d* = 1 between categories *HHA* and *HHH*).

To compare STDP rule sets of two simulations regardless of rule identity Fig. 5F-H, we computed the Wasserstein-1 distance (Earth Mover’s Distance,^89^) separately for *θ* = LTD/LTP and *γ* = *τ*_−_/*τ*_+_, using the log-transformed values of the three rules in each simulation. This is obtained by sorting the log-transformed values within each simulation and averaging the absolute differences between matched pairs:

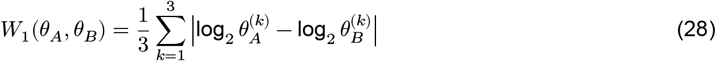

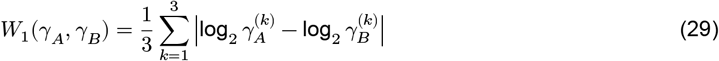

where *θ*^(*k*)^ and *γ*^(*k*)^ denote the *k*-th smallest values among the three STDP rules of a given simulation. We further min-max normalized each metric to the range [0, 1] across all pairs of parameter combinations.

To assess the significance of the correlations between JS divergence and rule distance (category distance, and the average LTD/LTP and *τ*_−_/*τ*_+_ distances; Fig. 5E-G), we used a permutation test. We randomly permuted the assignment of STDP parameters and rule-family identity across simulations, while keeping the sampled simulation pairs and their JS divergences fixed. For each of 5,000 permutations, we recomputed all pairwise distances and the corresponding Spearman correlation with JS divergence. Permutation p-values were calculated as the fraction of permuted correlations with an absolute value greater than or equal to the observed correlation.

### Comparison with data

To compare the motif distributions obtained above with the experimental data from ^31,34^ (Fig. 6), we considered the motifs in which each neuron is connected to at least one of the other two neurons. This excludes the disconnected motif and motifs involving only two-neuron interactions. We then computed absolute motif frequencies (i.e., without normalization relative to a random-graph null model) and normalized them to sum to one.

To quantify the similarity between the motif distribution of each simulation and each experimental dataset, we computed the Jensen-Shannon (JS) divergence as in Eq. (27), where *P* (m) is the simulated distribution, *Q*(m) is the experimental distribution, and 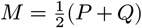. We classified simulations with JS divergence below 0.2 as having low divergence.

To characterize network activity in Fig. 6 D-F for the subset of simulations with low JS divergence from the mouse dataset^34^, which corresponds to simulations containing at least one Hebbian subpopulation, we simulated each network with plasticity switched off, i.e., all synaptic weights frozen at their final values, receiving the same external Poisson input (10 Hz). Simulations lasted approximately 300 s (the run length was set by the expected number of spikes given the final firing rates). Spike counts were binned in 50 ms windows for all activity measures.

We computed three measures: population firing rate, mean pairwise correlations, and dimensionality.

The population firing rate was computed from spike counts binned in 50 ms windows, summed across neurons and normalized by bin size and number of neurons, yielding an instantaneous population rate (Hz/neuron):

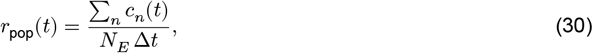

where *c*_*n*_(*t*) is the spike count of neuron *n* in bin *t*, Δ*t* = 50 ms, and *N*_*E*_ is the number of excitatory neurons. The population rate for a given simulation was defined as the time average of *r*_pop_(*t*).

Mean pairwise correlations were computed as the Pearson correlation coefficient between the binned spike counts of all pairs of excitatory neurons, averaged across pairs.

Population dimensionality was estimated from the spike-count covariance matrix of active excitatory neurons, computed from the binned spike counts. Denoting the eigenvalues of the covariance matrix as 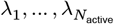, we used the participation ratio^61^:

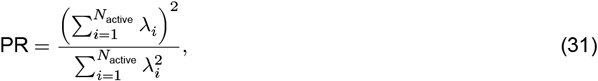

expressed as a percentage of *N*_active_. This measure quantifies the effective number of independent dimensions along which population activity varies, with PR → 100% indicating uniformly distributed variance across all neurons, and PR → 0% indicating variance concentrated in a single dominant mode.

To test for differences in population firing rate, mean pairwise correlations, and dimensionality across rule-family categories, we used the Kruskal–Wallis test and report the effect size as epsilon-squared. The three omnibus p-values were corrected for multiple comparisons across metrics using the Holm method. Significant omnibus effects were followed by Dunn’s post-hoc pairwise comparisons between categories, with Šidák correction for the number of pairs.

**Table 1:**
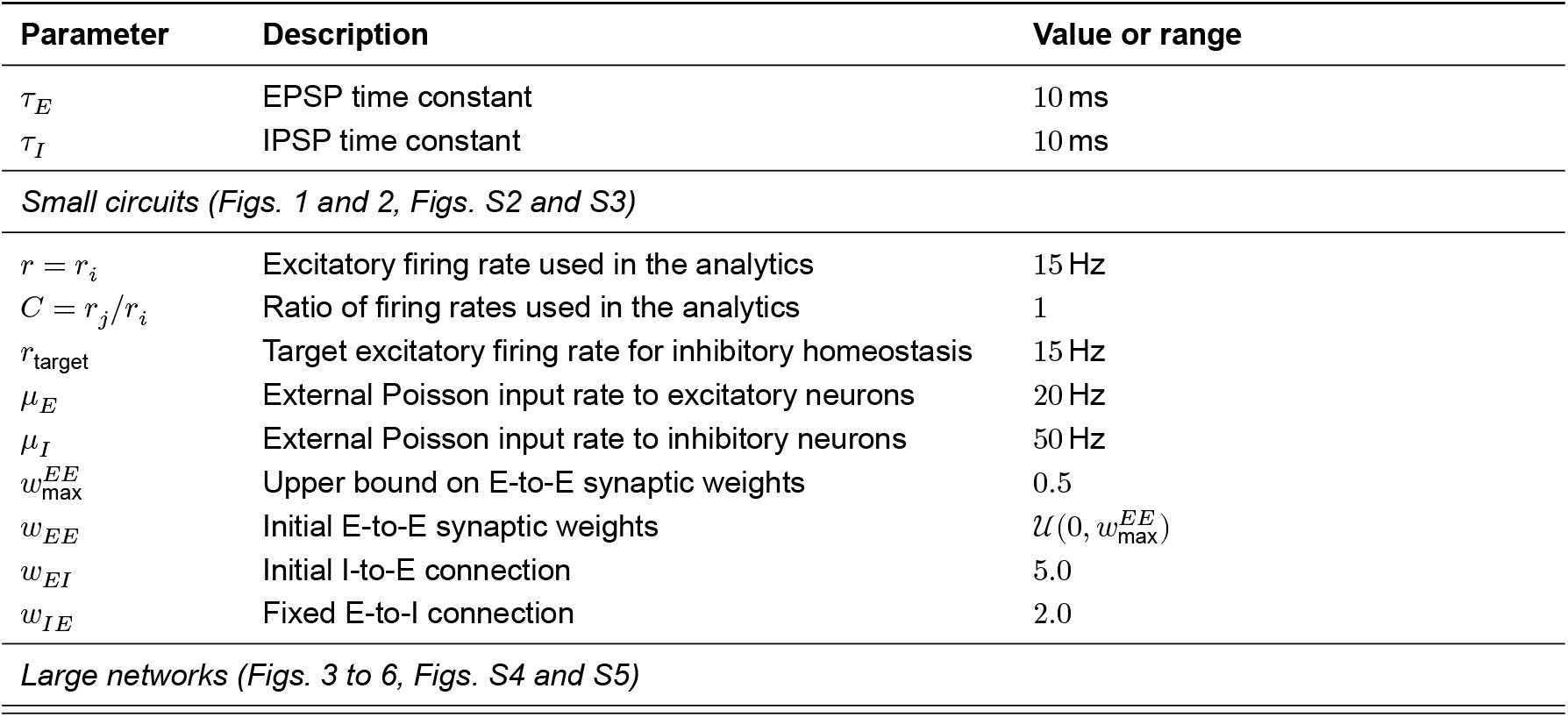
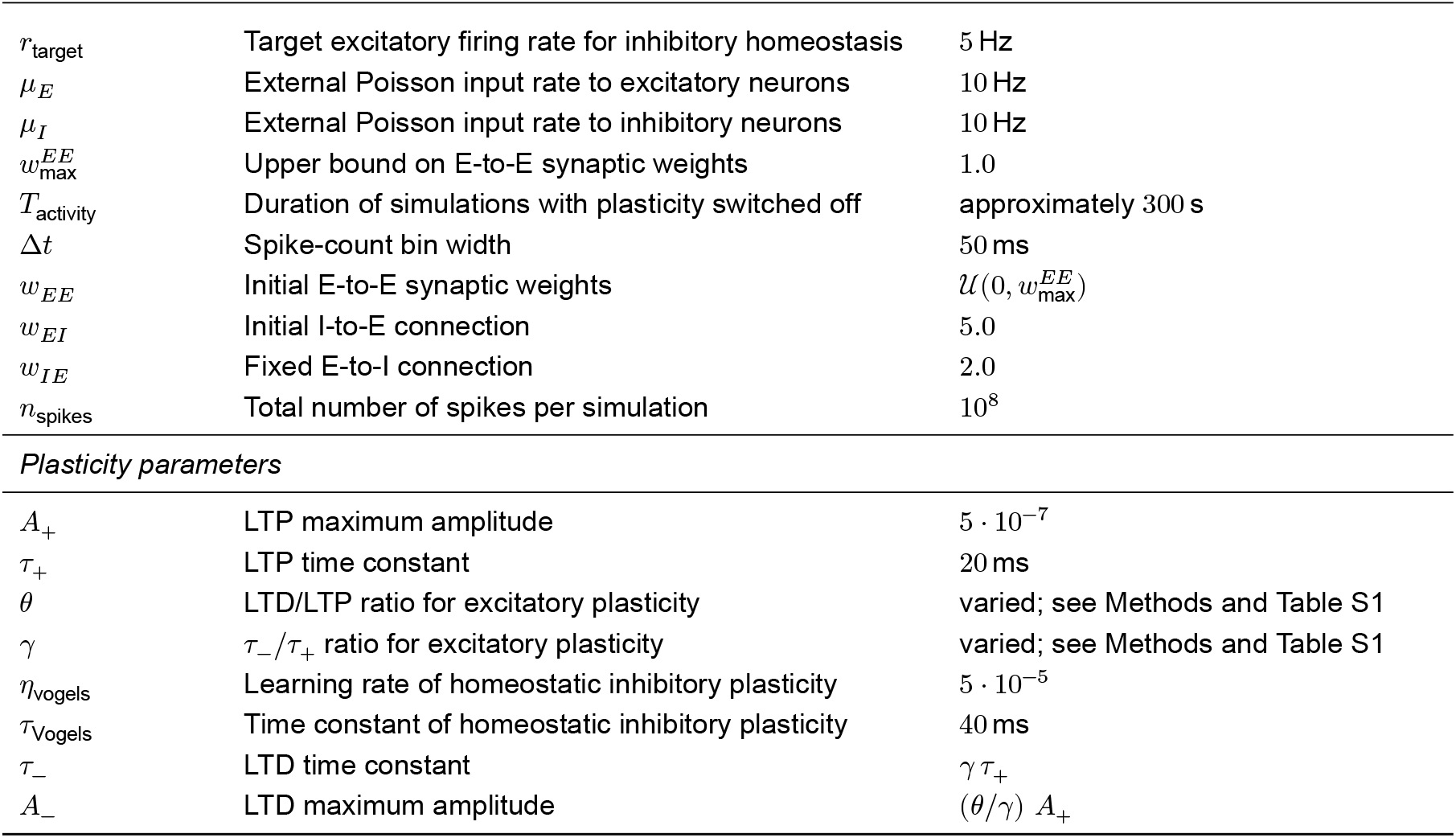
Model, simulation, and analysis parameters.

| Parameter | Description | Value or range |
|---|---|---|
| τ<sub>E</sub> | EPSP time constant | 10 ms |
| τ<sub>I</sub> | IPSP time constant | 10 ms |
| *Small circuits (Figs. 1 and 2, Figs. S2 and S3)* |  |  |
| r = r<sub>i</sub> | Excitatory firing rate used in the analytics | 15 Hz |
| C = r<sub>j</sub>/r<sub>i</sub> | Ratio of firing rates used in the analytics | 1 |
| r<sub>target</sub> | Target excitatory firing rate for inhibitory homeostasis | 15 Hz |
| μ<sub>E</sub> | External Poisson input rate to excitatory neurons | 20 Hz |
| μ<sub>I</sub> | External Poisson input rate to inhibitory neurons | 50 Hz |
| w<sub>max</sub><sup>EE</sup> | Upper bound on E-to-E synaptic weights | 0.5 |
| w<sub>EE</sub> | Initial E-to-E synaptic weights | 𝒰(0, w<sub>max</sub><sup>EE</sup>) |
| w<sub>EI</sub> | Initial I-to-E connection | 5.0 |
| w<sub>IE</sub> | Fixed E-to-I connection | 2.0 |
| *Large networks (Figs. 3 to 6, Figs. S4 and S5)* |  |  |
| r<sub>target</sub> | Target excitatory firing rate for inhibitory homeostasis | 5 Hz |
| μ<sub>E</sub> | External Poisson input rate to excitatory neurons | 10 Hz |
| μ<sub>I</sub> | External Poisson input rate to inhibitory neurons | 10 Hz |
| w<sub>max</sub><sup>EE</sup> | Upper bound on E-to-E synaptic weights | 1.0 |
| T<sub>activity</sub> | Duration of simulations with plasticity switched off | approximately 300 s |
| Δt | Spike-count bin width | 50 ms |
| w<sub>EE</sub> | Initial E-to-E synaptic weights | 𝒰(0, w<sub>max</sub><sup>EE</sup>) |
| w<sub>EI</sub> | Initial I-to-E connection | 5.0 |
| w<sub>IE</sub> | Fixed E-to-I connection | 2.0 |
| n<sub>spikes</sub> | Total number of spikes per simulation | 10<sup>8</sup> |
| *Plasticity parameters* |  |  |
| A<sub>+</sub> | LTP maximum amplitude | 5 · 10<sup>−7</sup> |
| τ<sub>+</sub> | LTP time constant | 20 ms |
| θ | LTD/LTP ratio for excitatory plasticity | varied; see Methods and Table S1 |
| γ | τ<sub>−</sub>/τ<sub>+</sub> ratio for excitatory plasticity | varied; see Methods and Table S1 |
| η<sub>vogels</sub> | Learning rate of homeostatic inhibitory plasticity | 5 · 10<sup>−5</sup> |
| τ<sub>Vogels</sub> | Time constant of homeostatic inhibitory plasticity | 40 ms |
| τ<sub>−</sub> | LTD time constant | γ τ<sub>+</sub> |
| A<sub>−</sub> | LTD maximum amplitude | (θ/γ) A<sub>+</sub> |

## Code availability

The code for our simulations is based on our Hawkes Simulator toolbox written in Julia (https://github.com/comp-neural-circuits/HawkesSimulator.jl) and is available at the following link: https://github.com/comp-neural-circuits/excitatory-stdp-motifs.

## Acknowledgments

This project has received funding from the European Research Council (ERC) under the European Union’s Horizon Europe research and innovation programme (ERC CoG FeedbackCircuits, grant agreement No. 101170267), and under the European Union’s Horizon 2020 research and innovation programme (ERC StG NeuroDevo, grant agreement No. 804824) and the Marie Skłodowska-Curie ITN SmartNets (grant agreement No. 860949).

Additional funding was provided by the Human Frontier Science Program Postdoctoral Fellowship (LT0005/2024-L). We thank all members of the “Computation in Neural Circuits” group, especially Joey Stawyskyj and Matthew P Getz for their useful discussions and insights.

## Supplementary Information

### S1 Analytic results

In the following sections we show the computation of the region boundaries in the parameter spaces shown in Fig. 2 of the main text.

### Asymmetric STDP rules

The asymmetric Hebbian (*L*_*H*_) and anti-Hebbian STDP (*L*_*A*_) rules are defined in Methods (see Eqs. (15) and (16)). Throughout, *A*_+_ and *τ*_+_ denote the amplitude and decay time constant of LTP, while *A*_−_ and *τ*_−_ denote those of LTD.

#### Hebbian STDP rules

For Hebbian STDP rules, the boundaries of the regions in the parameter space (Fig. 2A), are a result of the system described in Eq. (19) (Methods):

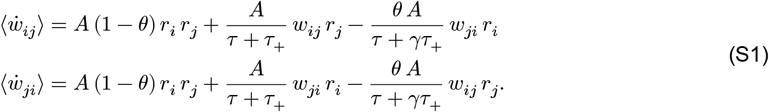

where *A* := *A*_+_*τ*_+_, *γ* := *τ*_−_/*τ*_+_, *θ* := (*A*_−_ *τ*_−_)/(*A*_+_ *τ*_+_). We imposed hard bounds on the synaptic weights:

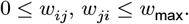

To further simplify the notation, we defined *r* := *r*_*i*_ and *C* := *r*_*j*_/*r*_*i*_.

We then found the following boundaries:

- A region of “unconnected motifs”, defined by:

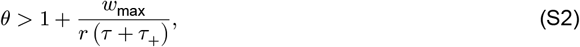

independent of *γ*. Here, the only attractor of the system is (0, 0), corresponding to loss of connection in both directions. Therefore, the steady-state triplet motif will be given by three unconnected neurons.

- A region of “all-to-all” connected motifs, defined by:

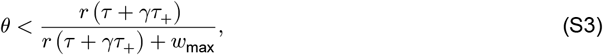

or, equivalently,

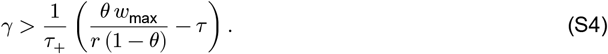

Here, the only attractor of the system is (*w*_max_, *w*_max_), corresponding to a bidirectional connection. Therefore, the steady-state triplet motif will be all-to-all connected. Intermediate cases involve:
- a region of “no reciprocal” connections, defined by 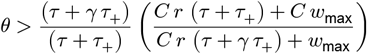, or, equivalently, 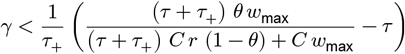
- a region of “no loss” of connections, defined by *θ* < 1,
- and a region of “only unidirectional”, given by the intersection of the two previous regions, where attractors are (*w*_max_, 0) or (0, *w*_max_), and the steady-state triplet motif can be either a feed-forward motif or a loop motif.

### Anti-Hebbian STDP rules

For anti-Hebbian STDP rules, we used the same parametrization as for the Hebbian rules, with *A*_+_ and *τ*_+_ denoting the amplitude and decay time constant of LTP, and *A*_−_ and *τ*_−_ those of LTD. Thus, we define *θ* := (*A*_−_ *τ*_−_)/(*A*_+_ *τ*_+_) and *γ* := *τ*_−_/*τ*_+_. For the anti-Hebbian rule, LTD occurs for positive time differences (*u* ≥ 0), whereas LTP occurs for negative time differences (*u* < 0). We found the following boundaries (shown in Fig. 2C). If *θ* > 1 and

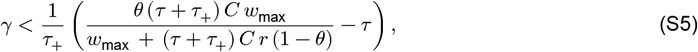

the only attractor of the system is (0, 0), corresponding to loss of connection in both directions. Therefore, the steady-state triplet motif consists of three unconnected neurons.

When *θ* > 1 and

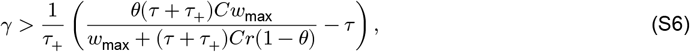

there are two possible types of attractors, namely (0, 0) and (*w*_max_, *w*_max_). In this case, the combination of these attractors facilitates the formation of new triplet motifs that could not be formed with Hebbian rules.

For *θ* < 1, the boundary is given by

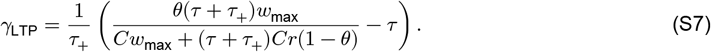

When *γ* > *γ*_LTP_, the only attractor is (*w*_max_, *w*_max_), corresponding to an all-to-all connected triplet motif.

Conversely, when *γ* < *γ*_LTP_, we found attractors with both connections within their allowed bounds.

### Symmetric rules

The symmetric STDP rule was defined as:

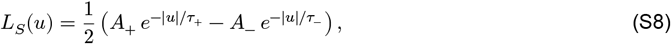

with *A*_+_, *A*_−_ > 0. We identified four regions of the parameter space depending on *θ* and the value

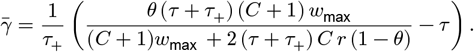

More precisely, we found the following regions:

- only bidirectional connections of the form (*w*_max_, *w*_max_), if *θ* < 1 and 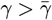;
- a line attractor, allowing unidirectional and bidirectional connections across a continuous range, if *θ* < 1 and 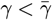;
- only loss of connections, if *θ* > 1 and 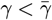;
- a combination of loss and bidirectional connections (*w*_max_, *w*_max_), if *θ* > 1 and 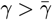.

Thus, the symmetric rule shares the attractors of the anti-Hebbian rule in three of the four regions, but also allows for a line-attractor region, where unidirectional connections are possible.

**Table S1:** Plasticity parameters of the example simulations.

| **Simulation** | **Rule type** | θ = A<sub>−</sub>τ<sub>−</sub>/A<sub>+</sub>τ<sub>+</sub> | γ = τ<sub>−</sub>/τ<sub>+</sub> | **Used in** |
|---|---|---|---|---|
| Example in Fig. 4 | Hebbian | 0.9 | 2.2 | Fig. 4C, D |
|  | Hebbian | 1.6 | 2.2 |  |
|  | Anti-Hebbian | 1.4 | 3.0 |  |
| *Examples in Fig. 5* |  |  |  |  |
| Example 1, sim 1 (HAA) | Hebbian | 1.64 | 2.73 | Fig. 5C |
|  | Anti-Hebbian | 2.98 | 2.56 |  |
|  | Anti-Hebbian | 0.90 | 1.77 |  |
| Example 1, sim 2 (ASS) | Anti-Hebbian | 0.78 | 1.89 | Fig. 5C |
|  | Symmetric | 1.15 | 0.91 |  |
|  | Symmetric | 1.01 | 0.87 |  |
| Example 2, sim 1 (HHH) | Hebbian | 2.27 | 0.47 | Fig. 5D |
|  | Hebbian | 0.97 | 2.05 |  |
|  | Hebbian | 1.78 | 2.98 |  |
| Example 2, sim 2 (HAS) | Hebbian | 1.78 | 1.81 | Fig. 5D |
|  | Anti-Hebbian | 0.47 | 2.28 |  |
|  | Symmetric | 0.49 | 2.40 |  |
| *Examples in Fig. 6* |  |  |  |  |
| Model 1 (HHA) | Hebbian | 2.83 | 0.40 | Fig. 6E, F |
|  | Hebbian | 2.77 | 1.49 |  |
|  | Anti-Hebbian | 1.17 | 1.53 |  |
| Model 2 (HHA) | Hebbian | 2.71 | 2.95 | Fig. 6E, F |
|  | Hebbian | 2.05 | 0.76 |  |
|  | Anti-Hebbian | 2.67 | 2.36 |  |
| *Examples in Fig. S4 & Fig. S5* |  |  |  |  |
| Switch between rules | LTD-Hebbian | 1.5 | 0.75 | Fig. S4 |
|  | LTP-Hebbian | 0.9 | 1.1 |  |
| Switch between rules | Hebbian | 1.0 | 1.0 | Fig. S5 |
|  | anti-Hebbian | 1.0 | 1.0 |  |

### S2 Supplementary figures

**Figure S1:**
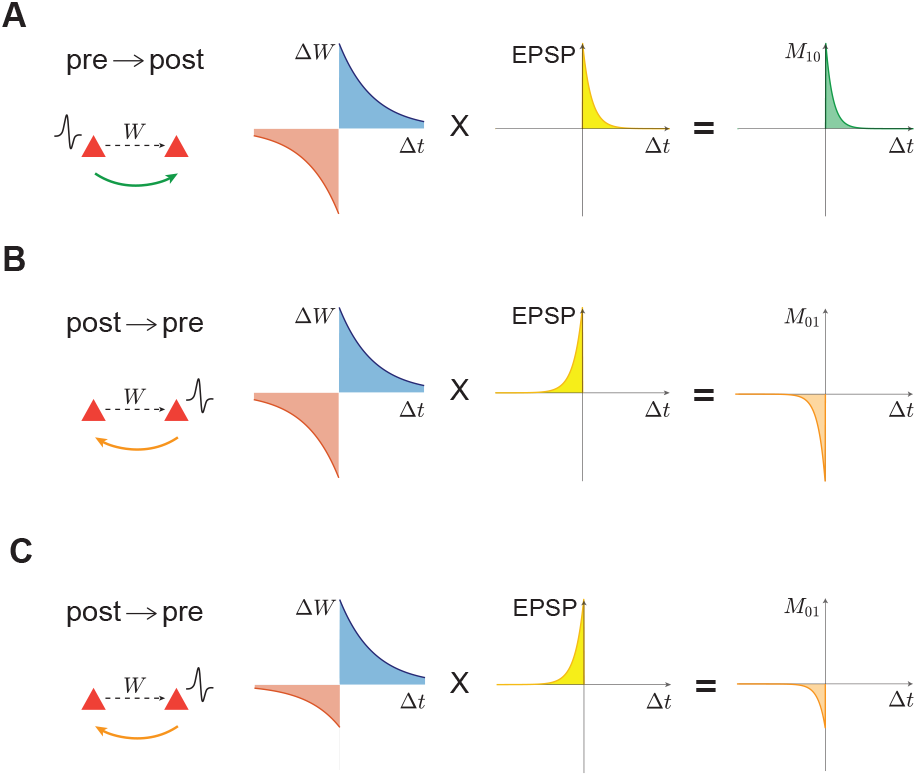
Motif coefficient calculation. **A**. Schematic illustration of Eq. 12 (see Methods): The term *M*_10_ (green integral on the right) represents the contribution to the synaptic change 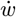 from presynaptic firing (left), and is calculated from the STDP kernel and the postsynaptic potential kernel (middle) (see also Montangie et al., 2020). **B**. Same as in **A**, but for *M*_01_, which describes the contribution from postsynaptic firing. **C**. Same as in **B**, but for an STDP rule with LTD/LTP < 1, resulting in a smaller magnitude of *M*_01_.

**Figure S2:**
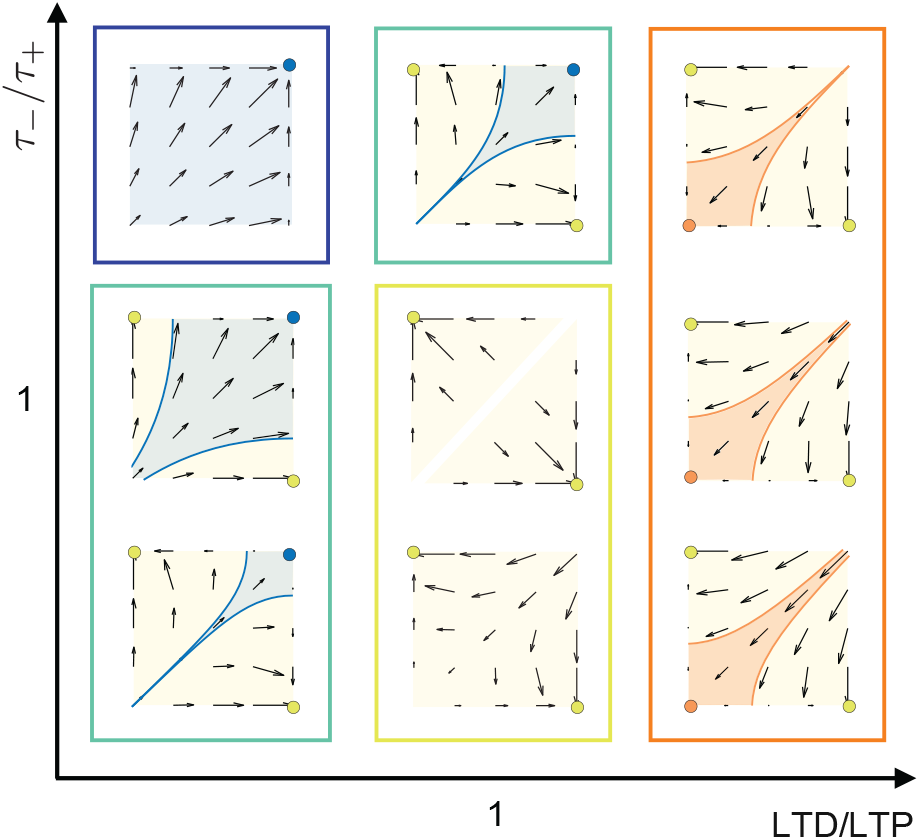
Phase planes for Hebbian rules. Phase planes for a given pair of connections evolving under single Hebbian rules, by varying LTD vs. LTP ratio and *τ*_−_/*τ*_+_. Blue, beige, and orange shaded areas denote different basins of attraction. Colored circles indicate the corresponding attractors. The colored boxes around (groups of) phase planes correspond to the emerging motif classes as described in the main text.

**Figure S3:**
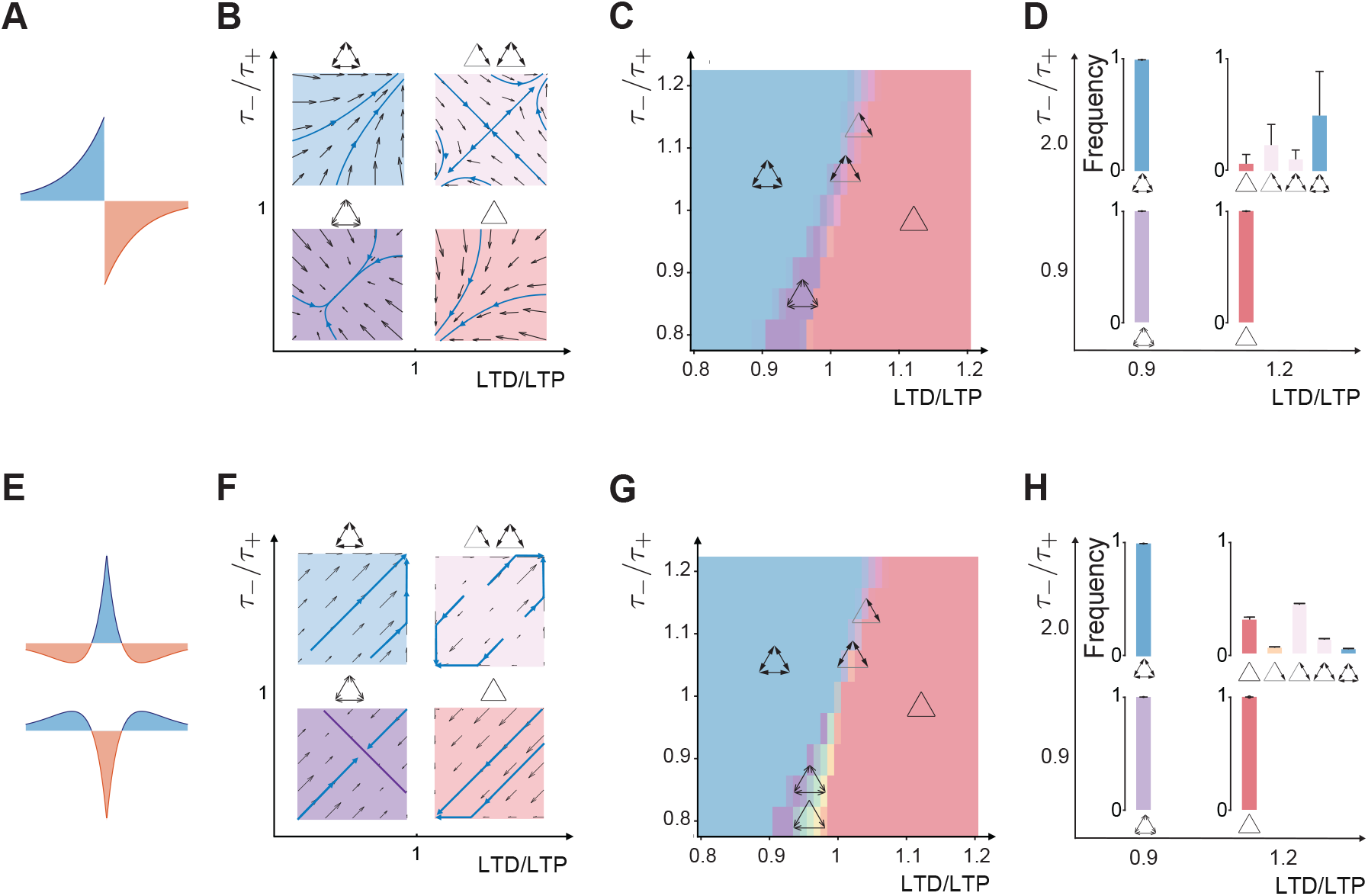
Anti-Hebbian and Symmetric rules - analytical prediction and network simulations. **A**. Exemplar anti-Hebbian rule. **B**. Phase planes for a given pair of connections evolving under single anti-Hebbian rules, by varying LTD vs. LTP ratio and *τ*_−_/*τ*_+_. Blue, pink, purple and red shaded areas denote different basins of attraction. Above each phase plane, we show all triplet motifs reachable from its dynamics by combining the three pairs. **C**. Motif classes obtained from simulations of three-neuron circuits with Hawkes dynamics under anti-Hebbian rules, as a function of LTD/LTP and *τ*_−_/*τ*_+_, averaged over 10 trials for each parameter combination. Colors represent classes of motifs: no connections (red), bidirectional/no connections (light purple), bidirectional with synaptic weights across a continuous range (dark purple), and only bidirectional (blue). **D**. Steady-state triplet motif distributions of simulated large networks (as in Fig. 3) in the excitatory population under anti-Hebbian rules, for different choices of LTD/LTP and *τ*_−_/*τ*_+_. Bar colors correspond to the region colors as in **C**. and Fig. 2 in the main text. Error bars denote mean ±standard deviation (*n* = 10 trials). **E**. - **H**. Analogous to **A-D**, but for the symmetric rules. The additional colors refer to a region where synaptic weights converge to a line attractor that includes unidirectional connections across a continuous range (purple phase plane in panel **F**, corresponding to LTD/LTP < 1 and *τ*_−_/*τ*_+_ < 1), which allows for the formation of additional motifs from the orange class (cf. main text.)

**Figure S4:**
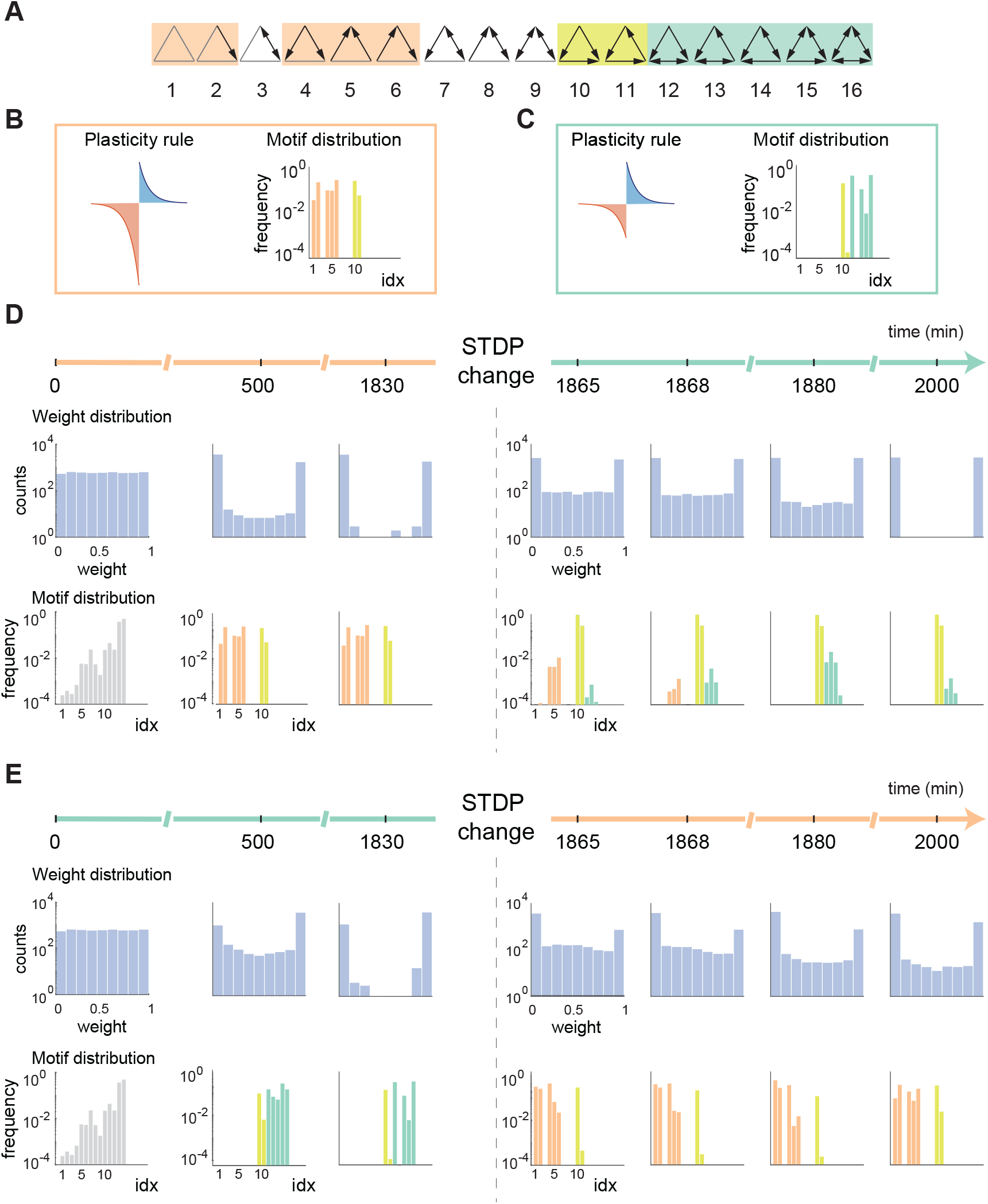
Time evolution from a LTD-dominated Hebbian rule to a LTP-dominated Hebbian rule and vice versa. (cf. Fig. S5). **A**. All sixteen triplet motifs. Colored boxes indicate classes of motifs (as in the main text) considered here. **B**. Schematic of a LTD-dominated STDP rule and corresponding motif distribution. Colors as in **A. C**. Schematic of a LTP-dominated STDP rule and corresponding motif distribution. Colors as in **A. D**. Evolution of weights (histograms on top row) and motif distributions (histograms on bottom row) over time, with STDP rule changing in the middle, from the LTD-dominated rule (panel **B**) to the LTP-dominated rule (panel **C**). **E**. Same as in **D**, but with the order of the rules reversed.

**Figure S5:**
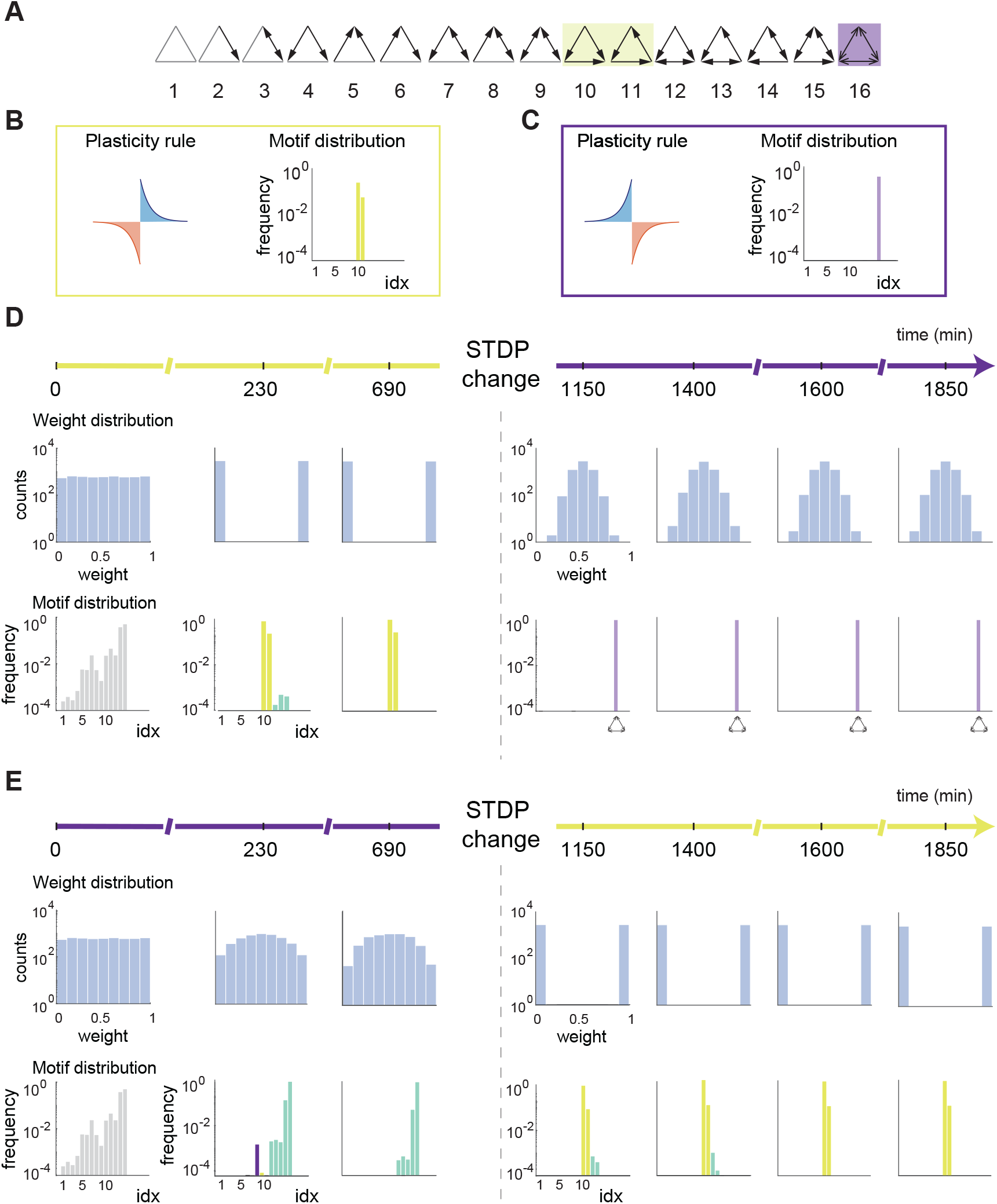
Time evolution from a Hebbian rule to an anti-Hebbian rule and vice versa. (cf. Fig. S4). **A**. All sixteen triplet motifs. Colored boxes indicate classes of motifs (as in the main text) considered here. **B**. Schematic of a Hebbian STDP rule and corresponding motif distribution. Colors as in **A. C**. Schematic of an anti-Hebbian STDP rule and corresponding motif distribution. Colors as in **A. D**. Evolution of weights (histograms on top row) and motif distributions (histograms on bottom row) over time, with STDP rule changing in the middle, from the Hebbian rule (panel **B**) to the anti-Hebbian rule (panel **C**). **E**. Same as in **D**, but with the order of the rules reversed.

